# Incorporating prior knowledge to seeds of adaptive sampling molecular dynamics simulations of ligand transport in enzymes with buried active sites

**DOI:** 10.1101/2023.09.21.558608

**Authors:** Dheeraj Kumar Sarkar, Bartlomiej Surpeta, Jan Brezovsky

**Affiliations:** Laboratory of Biomolecular Interactions and Transport, Department of Gene Expression, Institute of Molecular Biology and Biotechnology, Faculty of Biology, Adam Mickiewicz University, Uniwersytetu Poznanskiego 6, 61-614 Poznan, Poland; International Institute of Molecular and Cell Biology in Warsaw, Ks Trojdena 4, 02-109 Warsaw, Poland

**Author notes:** Corresponding author: JB.

**Keywords:** Protein-ligand, tunnels, ligand transport, seeding, adaptive sampling, kinetics

## Abstract

Given that most proteins have buried active sites, protein tunnels or channels play a crucial role in mitigating the transport of small molecules to the buried cavity for enzymatic catalysis. Tunnels can critically modulate the biological process of protein-ligand recognition. Various molecular dynamics methods have been developed for exploring and exploiting the protein-ligand conformational space to extract high-resolution details of the binding processes, one of the most recent represented by energetically unbiased high-throughput adaptive sampling simulations. The current study systematically contrasts the role of integrating prior knowledge while generating useful initial protein-ligand configurations, called seeds, for these simulations. Using a non-trivial system of haloalkane dehalogenase mutant with multiple transport tunnels leading to a deeply buried active site, these simulations were employed to derive kinetic models describing the process of association and dissociation of the substrate molecule. The more knowledge-based seed generation enabled high-throughput simulations that could more consistently capture the entire transport process, effectively explore the complex network of transport tunnels, and predict equilibrium dissociation constants, *k*_*off*_*/k*_*on*_, on the same order of magnitude as experimental measurements. Overall, the infusion of more knowledge into the initial seeds of adaptive sampling simulations could render analyses of transport mechanisms in enzymes more consistent even for very complex biomolecular systems, thereby promoting the rational design of enzymes with buried active sites and drug development efforts.

## 1. Introduction

Given the fact that molecular recognition is critical for all biological processes, in this context, the intrinsically dynamic and volatile nature of protein-ligand (un)binding processes makes it a long-standing quest to capture the high-resolution sampling and resolve meaningful kinetics of ligand binding processes in structure-based drug design^1,2^. Additionally, a ligand can prefer multiple routes of entry to interact with the environment of active site^3–5^. These routes, often referred to as tunnels, are seen to have equivalent importance as the catalytic properties of enzymes^6^. While in the majority of enzymes, the active site is buried^7,8^, the underlying molecular properties of the tunnels can control the entry and exit of ligands to a greater extent, specifically by gating residues^9^. In this context, the ligand binding processes via those transport pathways are a critical component in biocatalysis, also for identifying critical residues underlying the transport processes for mutagenesis and rational drug design^6,10^. Hence, protein tunnels are well-placed when considering improved catalysis and features like specificity and altered activity of small molecules. Because very often, tunnel lining residues or other gating residues can act as hot spots other than the active site residues^9,10^.

The transport processes, like a migration of ligands from the active site to the bulk solvent, are often connected with the requirement of overcoming a high energy barrier, resulting in the rare nature of such an event^11^. Because molecular dynamics (MD) simulations can observe biologically relevant processes even at atomistic resolution, they are extensively used to study mechanisms, dynamics, and functions of biomolecular complexes^12,13^. Numerous computational approaches have been developed in recent years to sample such rare events of ligand transport processes involving the association and dissociation of ligands and receptors^14^. These approaches benefit from the improvement of computational hardware in terms of GPUs as well as the implementation of various path sampling methods and methods for sampling rare events^15,16^. Specifically enhanced sampling methods like milestoning^17^, weighted ensemble^18^, Gaussian accelerated MD^19,20^, metadynamics^21,22^, adaptive sampling MD (ASMD) based on Markov state models (MSM)^5,23,24^, Random Acceleration MDs^5,25,26^, gained popularity in studying such rare events. While most methods use additional potential or force to bias the simulations along a designed collective variable, the ASMD methods utilizing MSMs can avoid such perturbations^27–29^. Extensive ASMD simulations have been successfully used to study ligand binding processes^5,24,30–33^. The ASMD is an energetically unbiased protocol comprising iterative rounds of intelligently respawned equilibrium simulations of protein-ligand configurations. This is achieved by using a scoring function to select the least explored configurations from a preliminary MSM build on the so far generated simulations and employing those configurations to initiate subsequent batches of simulations (called epochs)^28,29^.

Given the rising success of ASMD simulations in ligand transport studies, the impact of designing individual components in ASMD workflow on the efficacy of sampling relevant regions of protein-ligand configurational space is of interest^24,31,33–35^. Betz and Dror investigated the role of a scoring function for selecting the configuration for the successive iterations to partially overcome the exploration-exploitation tradeoff using the well-known test system of trypsin with benzamidine inhibitor and a more complex yet realistic system of membrane-bound adrenergic receptor β_2_ with dihydroalprenolol inhibitor^33^. They compared three scoring functions based on simple counts, in which states are resampled with probability inversely proportional to their occurrence in simulation; the population scores, which prefer states with smaller populations in MSMs; and hub scores, which select states with lower connectivity in MSMs, the measure of connectivity of states in MSMs. On the membrane-bound system, the count score could not govern the ASMD toward investigating inhibitor migration through the protein, focusing entirely on the membrane region. In contrast, the other two scores successfully sampled the relevant configurations. Hence, the use of more information-rich functions markedly benefited the study of ligand transport in more complex settings.

Here, we investigated the role of employing relevant information as early as in preparing initial seeding structures for ASMD. We designed four schemes (Figure 1A) from random positioning of the ligand around the protein to more knowledge-based poses of the ligand bound in the active site or along the tunnels precomputed from apo simulation. We tested the capabilities of ASMD, initiated from these seeding schemes, in exploring and exploiting the transport tunnels in haloalkane dehalogenase mutant LinB86 (Figure 1B), in which an additional functional tunnel was introduced *de novo*^10^. By performing intensive ASMD simulations of LinB86 with one of its substrates, 1,2-dibromoethane (DBE), for each scheme, we were able to compare to what degree the initial seeding impacts ability of ASMD to i) capture entire process of substrate association and dissociation, ii) identify metastable states adopted by substrate consistently, iii) predict kinetic parameters of the process, and finally iv) describe complexity of transport via multiple transport tunnels.

**Figure 1.**
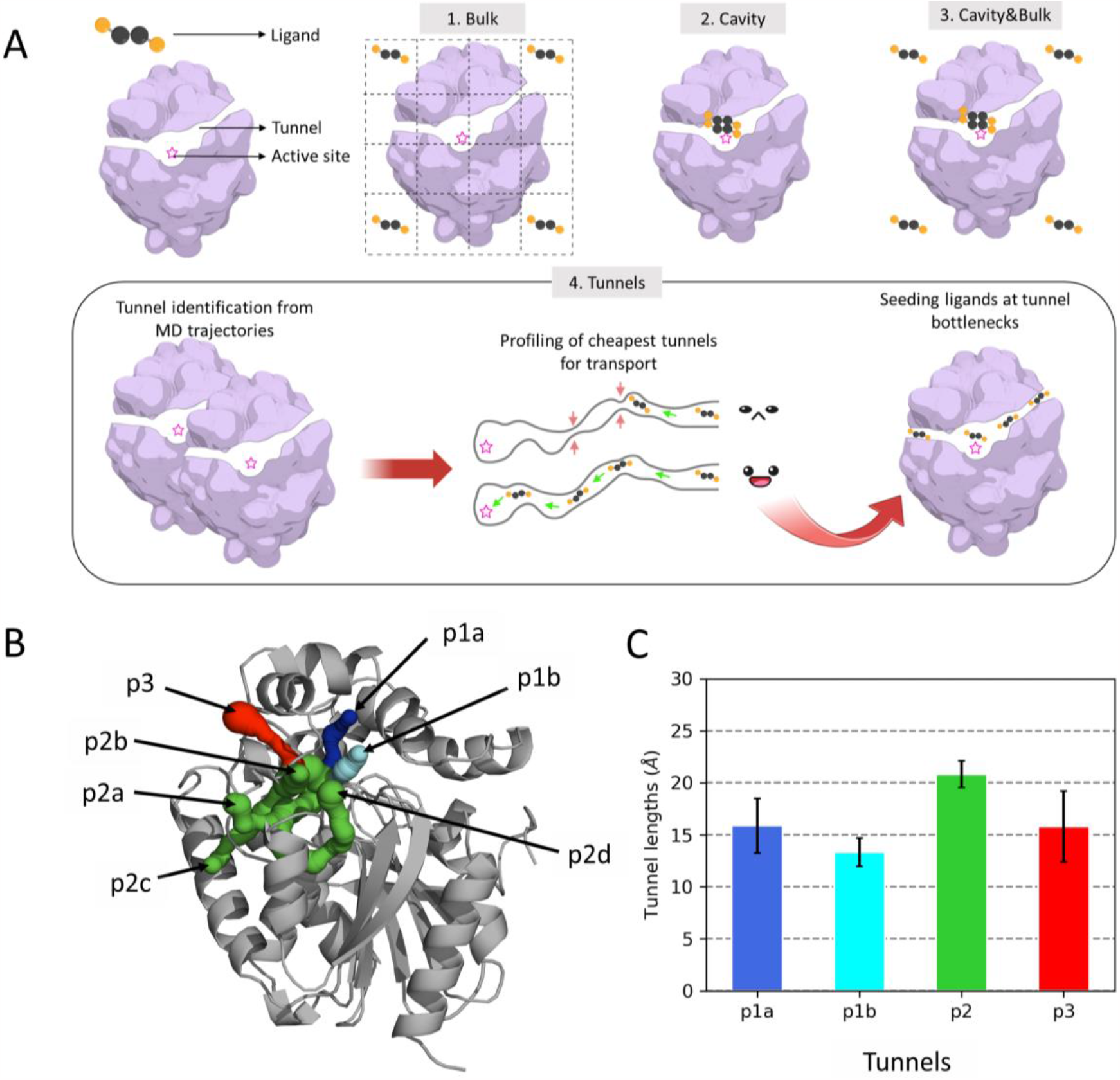
Overview of evaluated seeding schemes and model system used. A) Schematic representation of studied schemes and seeding of the substrate molecule from random to knowledge-based positions. B) Representative structure of tunnel network from 100 ns MD simulation of LinB86 (see Table S1 for other tunnel properties). The known tunnels are shown as sets of colored spheres: *p1a* (blue), *p1b* (cyan), four branches of *p2* (green), and *p3* (red). The protein structure is shown as a gray cartoon. C) Average lengths of ensembles of the known tunnels observed in MD simulation.

## 2. Materials and Methods

### 2.1. Seed generation for ASMD simulations

The input model was based on the crystallographic structure of the mutant of haloalkane dehalogenase enzyme LinB86 (PDB code: 5LKA). The protein structure was further protonated using H++ web server^36,37^ pH 8.5. The protein was modeled using the AMBER ff14SB^36^ force field and the substrate DBE with the General Amber Force Field -GAFF^38,39^. The partial atomic charges on the DBE molecule were derived using multi-conformational, multi-orientational restrained electrostatic potential fit^40^. Each DBE conformation was geometry optimized at MP2/6-31G(d) level of theory, and their multi-orientational molecular electrostatic potential was calculated at HF/6-31G(d) level using Gaussian v09^41^. Finally, two-stage charge fitting was conducted for all conformations and orientations using *resp* and *antechamber* modules of AMBERTools18^42^.

The substrate molecule was placed according to the four designed schemes to investigate the role of knowledge in ASMD seeding systematically (Figure 1A). DBE was placed at 30 different positions in each scheme (Figure S1) as follows. i) In the *Bulk* scheme, DBE was positioned on an equally spaced grid in the bulk solvent surrounding the protein using *“drawgridbox [selection], nx=5, ny=5, nz=5, padding=5, lw=1, r=0, g=0”* function of PyMOL^43^. ii) In the *Cavity* scheme, DBE was docked to the enzyme’s active site using AutoDock Vina^44^. The 30 docked poses were derived by defining the grid box centered at COM of catalytic residues (N38, D108, W109, and H272) with a dimension of 22.5 Å and exhaustiveness of 1000. iii) In *the Cavity&Bulk* scheme, 15 DBE positions were taken from the *Cavity* scheme and 15 DBE positions from the *Bulk* scheme. Finally, iv) in the *Tunnels* scheme, putative transport tunnels in LinB86 were detected from 100 ns trajectory of ligand-free LinB86 simulation, and the most open tunnels were then explored for binding of DBE molecules along these tunnels. Finally, the composite tunnels, formed from parts of tunnels with conformations ensuring minimal energy costs for DBE migration, were generated (see Text S1 and Figures S2-S9 for details of this protocol).

The generated protein-ligand complexes were then solvated based on the 3D Reference Interaction Site Model theory^45^ using the Placevent^46^ algorithm. Such a system was then processed with the *tleap* module of AMBERTools18, placing the pre-solvated proteins in the octahedral box of OPC water molecules^47^ to the distance of 10 Å and neutralizing them with counter ions (Na^+^ and Cl^-^) to the ionic strength of 0.1 M. Finally, the hydrogen mass repartitioning method was applied to produce topologies to enable 4 fs timestep^48^.

### 2.1. Equilibration MD simulations of seeds

The systems were then minimized and equilibrated using PMEMD and PMEMD.CUDA modules^49^ of AMBER18^42^, respectively. All complexes were energy minimized in five consecutive stages, each composed of 100 steps of the steepest descent followed by 400 steps of the conjugate gradient method, with gradually decreasing restraints on the protein atoms (initially 500 to heavy atom, and later restraints of 500, 125, 25, and 0.001 kcal.mol^-1^.Å^-2^ applied only to the backbone atoms). Minimization was followed by 20 ps heating from 0 to 200 K in the NVT ensemble using the Langevin thermostat with a collision frequency of 2 ps^-1^ and coupling constant of 1 ps while keeping the protein heavy atoms restrained with a force constant of 5 kcal·mol^-1^·Å^-2^. Next, the temperature was raised to the target value of 310 K in 100 ps of NVT simulation and kept constant for 900 ps, employing the same parameters as previously described. This was followed by NPT simulation at 1 atm enforced by the weak-coupling barostat with a coupling constant of 1 ps using positional restraints of 5 kcal·mol^-1^·Å^-2^ on the backbone atoms for 1 ns, followed by 1 ns without any positional restraints. All MD simulation stages were run using a 4 fs timestep enabled by SHAKE^50^ and hydrogen mass repartitioning algorithms, periodic boundary conditions, and particle mesh Ewald method^51^. The trajectories were generated by saving coordinates every 20 ps. The MD trajectories were analyzed using the *cpptraj* module of AMBERTools23^52,53^. The last snapshots from the unrestrained simulation were used as the initial input structures for ASMD.

### 2.2. High throughput ASMD to study substrate un(binding) processes

The ASMD was set up with 30 epochs, each consisting of 30 separate production simulations. To build an MSM model after each epoch, we used the distances between the Cα atoms of the protein and four heavy atoms of DBE and reduced the high dimensional space to three dimensions using time-dependent component analysis (TICA)^54^ with a lag time of 2 ns. The ASMD simulations were performed using HTMD v1.13.10^27^ and AMBER18^42^ software packages. The equilibration phase in HTMD consisted of two 250 ps NVT and NPT simulations, during which the systems were heated from 0 to 310 K with a Langevin thermostat and harmonic positional restraints to the backbone atoms with a force constant of 5 kcal·mol^-1^·Å^-2^. Finally, a 50 ns unrestrained production MD was performed in the NVT ensemble using a weak-coupling thermostat and a saving frequency of 100 ps. Such ASMD runs were performed in three replicates for each investigated seeding scheme.

### 2.3. Final MSM construction and validation

All the MSM were built using the HTMD^27^, which internally uses PyEMMA program^55^, following the standard PyEMMA protocol. The high dimensional data from adaptive sampling projected with distance feature was reduced to three dimensions using TICA^54^ with a lag time of 2 ns. Next, the reduced TICA coordinates were clustered into 1000 microstates using the MiniBatchKMeans^56^ method. The metastable states were lumped using PCCA++ method^57^, with the number of metastable states based on spectral analysis^55^ and verified against plots of linear implied timescales (Figures S10-S12). The lag time of 20 ns was used during MSM construction. Finally, the Chapman-Kolmogorov^58^ test was performed to confirm the Markovianity of the generated MSM models (Figures S13-S15).

### 2.4. MSM analysis and comparison

In order to quantify the ability of ASMD to sample the whole (un)binding process of DBE, the distances between the center of mass (COM) of the DBE molecule and COM of three catalytic residues (N38, D108, and W109, Figure S16A) were measured from the ~900 trajectories for each replicate using *cpptraj* module of AMBERTools23^52,53^. Based on this distance, we can define the location of DBE in the active site (0-5 Å), tunnel (5-19 Å), and bulk (>19 Å). The cutoff of 19 Å for tunnels was derived from the average lengths of investigated tunnels measured by CAVER (Figure 1C and Table S1). Finally, the transition path theory approach implemented in PyEMMA was used to derive transition probability matrices and compute the mean first pass times of each association and dissociation process in MSMs. Here, the metastable states with the most prevalent bound conformation of DBE were used as sink states, while the metastable states featuring DBE mainly in the bulk solvent were considered as source states to perform the transition flux analysis and derive the transition probabilities and kinetics rates. Furthermore, the most frequently occurring bottleneck residues were shortlisted from the CAVER results as follows (Figure S16B-E): *p1a* (D147, F151, and V173), *p1b* (D147, W177, and L248), *p2* (L211 and L248), and *p3* (L143, F151, and I213) and the distance between their COM to COM of DBE was calculated to assess localization of DBE with respect to these tunnels.

Ensembles of 1000 representative structures of metastable states generated from individual MSMs were clustered to establish the correspondence of these metastable states across explored schemes. For this purpose, mean, 25^th^, 50^th^, and 75^th^ percentiles were calculated for each set of characteristic distances to bottleneck residues and catalytic machinery described above (Figure S16). They were used cumulatively as a vector of 20 variables describing each metastable state. Principal component analysis (PCA) implemented in the Python scikit-learn library^56^ was used to reduce the dimensionality of each vector. The set of the first three principal components for each metastable state was clustered with HDBSCAN^59^ using *min_cluster_size* of 2, with the remaining parameters kept as default.

### 2.5. Analysis of substrate utilization of tunnels

Time-evolution of distances (Figure S16) for the entire set of trajectories was used to estimate the approximate position of the ligand in the context of the tunnel network. By tracking the change of the relative position, the movement through a particular tunnel was assigned where possible. Therefore, the approximate tunnels’ utilization was estimated across investigated schemes by analyzing the transition between subsequent positions. The procedure was composed of three stages as follows.

#### i) Position assignment

First, the closest bottleneck at a particular frame to the DBE molecule was defined. Further, this information was used to define the approximate length of the closest tunnel, i.e., the distance between the COM of catalytic machinery and the COM of the particular bottleneck. These two distances were contrasted with the distance of the ligand to the catalytic machinery, which altogether resulted in the identification of the approximate ligand position. Importantly, at this point, additional parameters were introduced to classify the ligand position, namely *bt_cutoff_along=2*.*0 Å* defining the region around the bottleneck, distinguishing whether the ligand is in the bulk, bottleneck *i)* region or tunnel, and *bt_cutoff_across=5*.*0 Å* that defines whether the ligand is not too far from the bottleneck horizontally in case it is within the bottleneck region. Considering these three distances and introduced cutoffs, the following scenarios and corresponding ligand states were considered:

- *Bulk (out_)*: Ligand is further from the active site than the sum of tunnel length and *bt_cutoff_along*.
- *Bottleneck (bt_)*: Ligand is within the bottleneck region, either further than the tunnel length or closer than the tunnel length but within *bt_cutoff_along* and *bt_cutoff_across*.
- *Unknown bottleneck (bt_unknown)*: Ligand is within the bottleneck region, either further than the tunnel length or closer than the tunnel length within *bt_cutoff_along* but exceeding the *bt_cutoff_across*.
- *Inside (in_)*: Ligand distance to catalytic machinery is shorter than the tunnel length decreased by the *bt_cutoff_along*.

#### ii) Transition detection and classification

Considering defined states for each frame, the transitions between bulk (out) and interior (in) and vice versa were identified. Transitions via bottleneck regions (in-bt-out or out-bt-in) were also considered. In the case that the mismatch between assigned tunnel in-out/out-in was detected, we applied an additional *dist_tolerance=1*.*0 Å* parameter, which defined the tolerance distance that is considered for swapping the classification of one of the sides of the transition, promoting the tunnel that was seen in the bottleneck region for scenarios where the intermediate state was seen. The transitions were tracked as follows:

- If the transition occurred from the bulk to inside or from the inside to bulk directly -the transition in-out/out-in was assigned by applying the *dist_tolerance* for cases where the mismatch between both sides occurred.
- If the ligand moved from the interior to the bottleneck region or from the bulk to the bottleneck region – the transition was not assigned, only the information regarding the temporary state.
- If the temporary state was a bottleneck and the closest tunnel changed, the transition was not assigned; only the bottleneck temporary state was updated.
- If the temporary state was a bottleneck and the ligand moved to the same general state but related to a different tunnel, the transition was not assigned; only the general state was updated.
- If the temporary state was a bottleneck and the ligand moved to the other general state (from in to out or from out to in), the transition was assigned, also collecting the information about the bottleneck used applying the *dist_tolerance* for cases where the mismatch between both sides occurred promoting the tunnel of the assigned transition bottleneck state.

#### iii) Characterization of tunnel utilization

Finally, all types of unique transitions were counted across all simulations from each scheme and averaged across three replicates performed for each scheme. Importantly, we applied the following classification to assign transitions to particular categories:

- Tunnel (*p1a, p1b, p2*, and *p3*) – all transitions that passed through the bottleneck of a particular tunnel or the direct transitions in-out or out-in related to the same tunnel on both sides;
- Mixed – all direct transitions in-out or out-in, where both sides of transitions differ even considering applied distance tolerance;
- Unknown – all transitions that crossed through the unknown bottleneck.

## 3. Results and Discussions

Overall, ~900 MD trajectories (450,000 frames) with aggregated simulation time of 45 µs were produced using ASMD of LinB86-DBE complexes generated according to all four studied schemes (Table S2). For each scheme, ASMD was performed in three replicates to evaluate the abilities to consistently describe the transport processes in its entirety, focusing on the convergence among the ASMD replicates, the degree of quantitative agreement with experimental data, and the ability to consider the transport via all known tunnels.

### 3.1 Capturing DBE association and dissociation processes in LinB86

In order to study the applicability of the studied schemes, we initially investigated how effectively each scheme could sample the endpoints of the processes, i.e., the bound and unbound states of DBE in the active site cavity of LinB86 and bulk solvent, respectively. Those states could be effectively defined by the distance of DBE COM from the COM of three catalytic residues located at the bottom of the cavity (Figure S16A), defining the bound states within 5 Å distance, while the unbound state samples primarily distances above 19 Å, which are further than the length of the longest tunnels present in LinB86 (Figure 1C). The DBE explored the unbound state in all schemes and replicas for a substantial fraction of cumulative ASMD trajectories (Figure 2). Even in the *Cavity* scheme initiated from the DBE molecule bound deep in the active site, the substrate reached the bulk solvent, generating a minimum of 12 % unbound states. Over 10,000 unbound states were generated after at most seven epochs of ASMD simulations (Figure 3). Foreseeably, the unbound states were most prevalent (> 31 %) in the simulation seeded with DBE placed in the bulk solvent around the enzyme (scheme *Bulk*).

**Figure 2.**
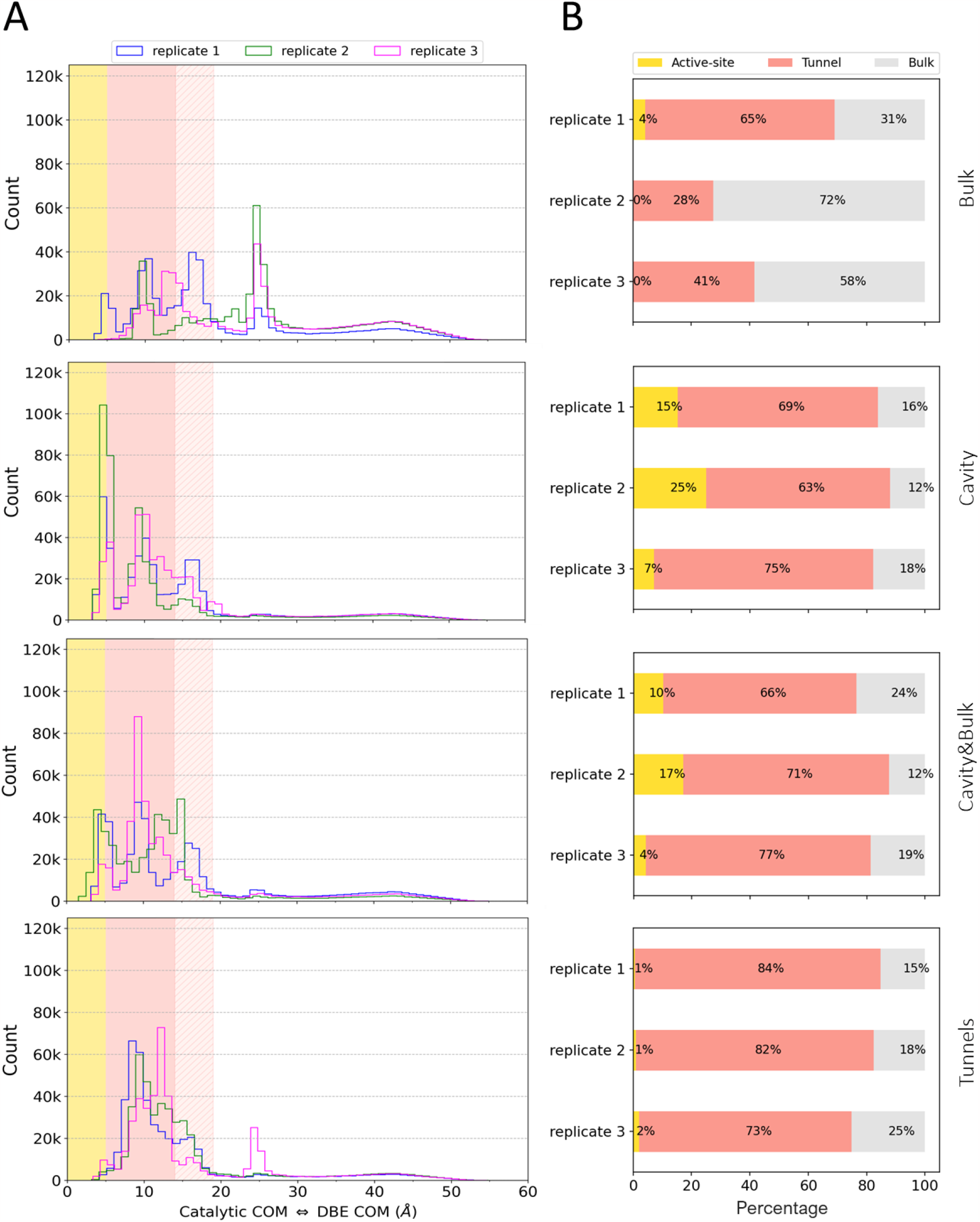
Substrate (un)binding to the active site of LinB86 captured by ASMD simulations with four seeding schemes. A) The distance distribution of DBE to catalytic residues for three replicated ASMD for each seeding scheme. The regions corresponding to DBE in the active site (0-5 Å), shortest (p1b, 5-14 Å) and longest (p2, 14-19 Å) tunnel lengths (Figure 1C), and bulk solvent (>19 Å) are highlighted as gold, pink, shaded pink, and white, respectively. The distances are between COM of DBE and COM of catalytic residues (N38, D108, and W109), measured in 45 µs simulations. B) The fraction of DBE seen in individual regions.

**Figure 3.**
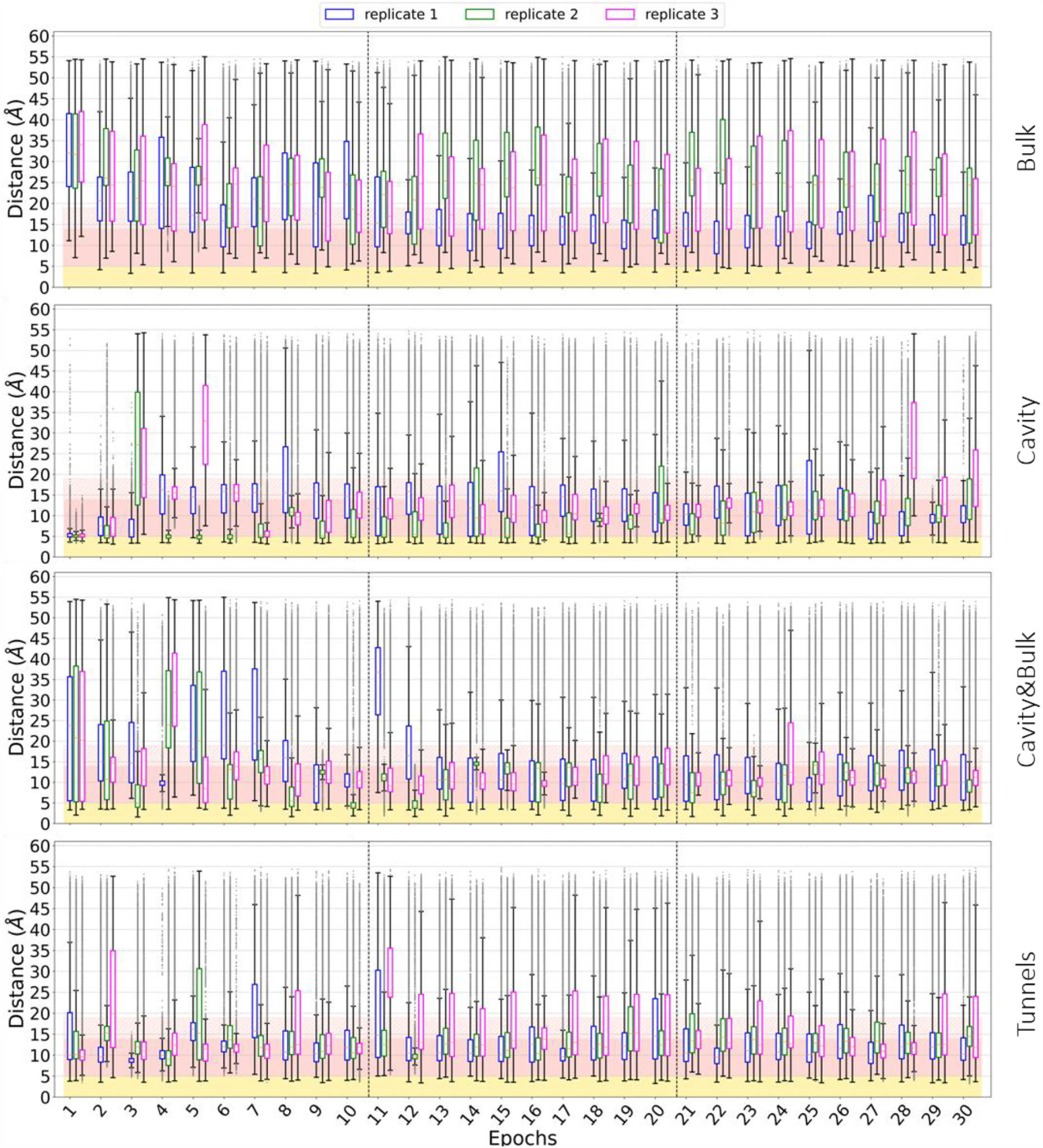
An epoch-wise sampling of the distance of DBE to catalytic residues of LinB86. The regions corresponding to DBE in the active site (0-5 Å), shortest (p1b, 5-14 Å) and longest (p2, 14-19 Å) tunnel lengths (Figure 1C), and bulk solvent (>19 Å) are highlighted as gold, pink, shaded pink, and white, respectively.

Concerning the ability of ASMDs to reach the bound pose of DBE in the buried active site of LinB86, all schemes except for *Bulk* could consistently sample the bound states in all three replicates. In the case of the *Bulk* scheme, the DBE molecule was able to find a path to the active site in replicate 1, producing a total of 4 % of simulations in the bound state (Figure 2), with a significant ensemble of more than 1,000 bound configurations sampled already until the fifth epoch (Figure 3). However, no bound state was observed in the other two replicated ASMD from the *Bulk* scheme (Figures 2 and 3). This is no surprise since unbiased simulations of ligand associations are generally rather time-consuming, even for less complex systems^60–62^.

Among the remaining schemes, *Cavity* AMSDs exploited the bound states the most frequently, as expected from the initial seeding with docked poses of DBE (Figure 2). Such setup led to the accumulation of over 5,000 bound states already during the first epoch in all three replicates (Figure 3). Such behavior was also partially retained in the *Cavity&Bulk* scheme, where more than 2,000 bound states were systematically observed in the first epoch of ASMDs. Here, the additional seeds of DBE placed in the bulk solvent resulted in considerable sampling of more than 10,000 unbound states within the first three epochs of ASMDs, about twice faster than in the pure *Cavity* scheme (Figure 3). Finally, we have observed the DBE spending most of the time exploring the regions corresponding to transport tunnels in ASMDs from the *Tunnels* scheme (Figure 2). Having sufficient coverage of bound and unbound states, we progressed to the creation of MSMs from the assembled trajectories and the calculation of kinetic parameters of descriptions of (un)binding processes. Due to the lack of bound states in the *Bulk* scheme, these AMSDs were not considered for constructing MSMs.

### 3.2. Identifying metastable states of DBE interacting with LinB86 and predicting kinetic parameters from MSMs

To further test the capabilities of the studied seeding schemes in the diversity and consistency of identified metastable states, we have generated MSMs from the individual ASMD replicates. These MSMs consisted of three to six metastable states for the *Cavity* (Figures S17-S19) and *Cavity&Bulk* (Figures S20-S22) schemes, whereas six to eight metastable states were identified in MSMs from the *Tunnel* schemes (Figures S23-S25). To understand the mutual correspondence among these states across all generated MSMs, we have generated 1,000 representative structures of each metastable state and measured the distances of DBE to the catalytic residues, as well as to the bottlenecks of the known transport tunnels in LinB86 (Figure S16). These distances represent fingerprints characterizing the metastable states (Figures S26-S28), clearly identifying not only unbound and bound states but also their alignment to individual transport tunnels.

Finally, those unified fingerprints enabled us to cluster the metastable states (Figure S29), forming the unified non-redundant ligand states (*ULS*) across all MSMs (Figure 4A). The only state consistently present in all replicates of each seeding scheme (Figure 4B) was *ULS1*, which corresponded to the DBE molecules in the bulk solvent. *ULS2-ULS5* all represented DBE molecules inside the catalytic cavity, with DBE bound closest to the catalytic residues in *ULS2*, which was found only in the MSMs of the *Cavity* scheme. In *ULS3* the substrate was placed closer to the cavity center, while in *ULS4* and *ULS5*, the substrate was located near the exit from the cavity in the direction of *p1* or *p3* tunnels. *ULS6-ULS9* featured the DBE molecule bound on the LinB86 surface at the entrances to *p3, p2a, p2c*, and *p2d* tunnels, with the *p3* tunnel entrance (*ULS6*) being the most prevalent across the MSMs (Figure 4B). Curiously, in replicate 2 from the *Cavity&Bulk* scheme, we have observed several metastable states forming *ULS10*, which were composed of the DBE molecules exploring the cryptic pocket located back-to-back with the canonical active site cavity of LinB86 with the entrance located on the opposite side of the enzyme structure with respect to the *p1* tunnel entrance.

**Figure 4.**
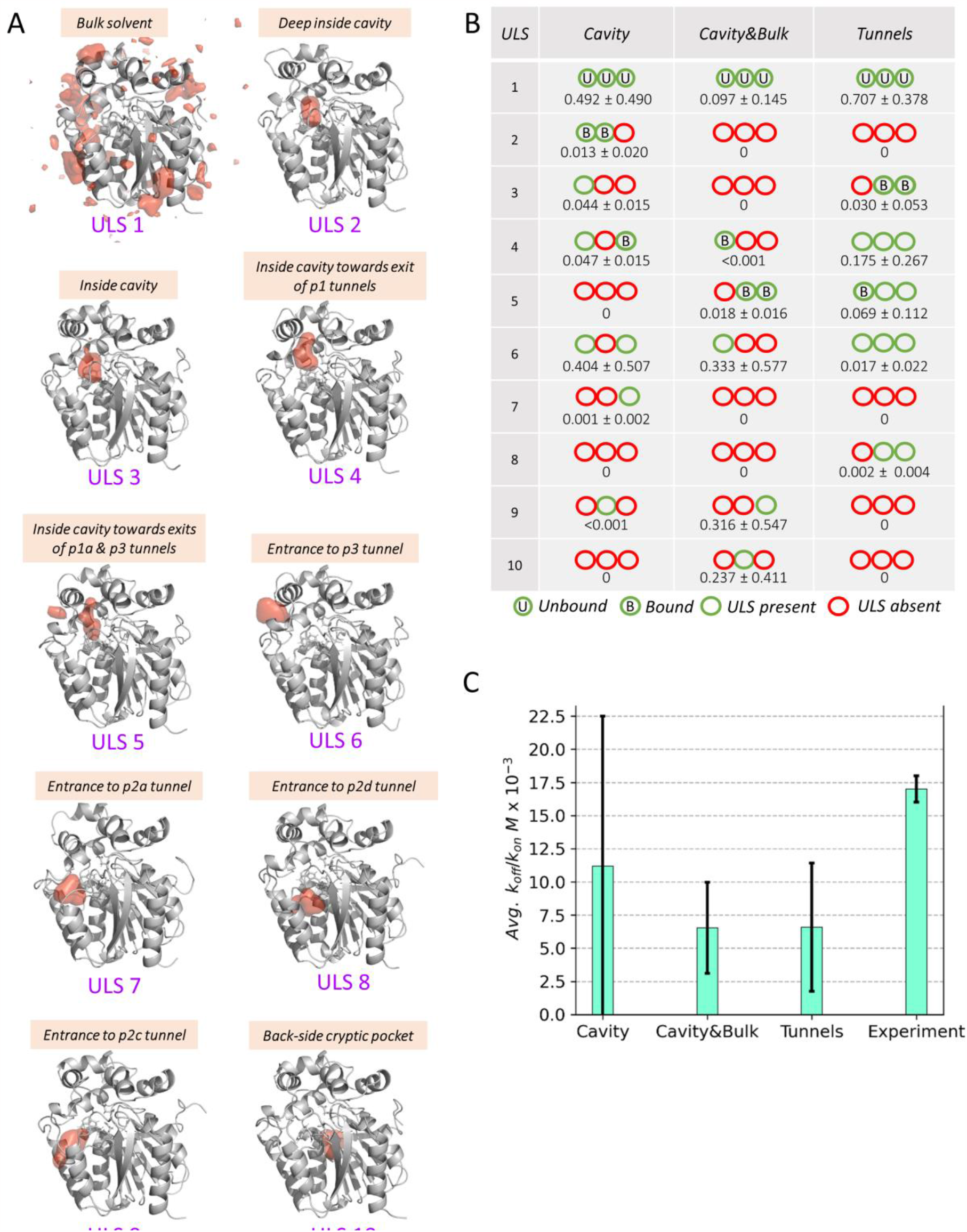
Inference into the (un)binding process of DBE to LinB86 from MSM analysis. A) Structurally unified ligand states (ULS) identified among all metastable states (Figures S17-S25) resolved by MSM analysis of three replicated ASMD simulations initiated from the studied seeding schemes. Protein structure is shown as a gray cartoon while the region occupied by DBE molecule in 20 % (1 % for bulk solvent state) of 1,000 structures representing given ULS is shown as red surface. B) The presence of ULS among metastable states in each ASMD replicates with their average probabilities. The unbound and bound metastable states used as source and sink states during the mean first pass time analyses are highlighted. C) Average equilibrium dissociation constants derived from MSMs as a ratio of dissociation and association rates (Figure S30). The experimental *k*_d_ was obtained from ^66^. The data represents mean±stdev from the three replicates.

Some identified *ULSs* were also observed in the recent study of transient binding sites on the LinB wild-type conducted with seven halogenated compounds, including DBE molecules^63^. Out of nine sites, three could be matched to *ULSs* as follows: i) *site 5* corresponded to *ULS6*, the entrance to the *p3* tunnel, ii) *site 9* overlayed with *ULS9*, the entrance to *p2c* tunnel, and iii) *site 4* aligned to *ULS8*, the entrance to *p2d* tunnel. Such agreement suggests conservation of those interaction sites between LinB wild-type and LinB86 mutant despite the substitutions introduced into the *p1* and *p3* tunnels of the mutant. Considering the identification of *ULS* in replicated MSMs, the *Tunnels* scheme exhibited the best consistency since four *ULS* were found in all three replicates, while the other two *ULS* were found in two replicates. In contrast, only unbound *ULS1* was systematically found in the *Cavity* and *Cavity&Bulk* schemes. In fact, those two schemes frequently led to the formation of singleton *ULSs*, present in one MSM replicate only.

Finally, we have calculated the equilibrium dissociation constant (*k*_*d*_) from the rates of DBE association and dissociation predicted from MSMs by the mean first pass time analyses. In contrast with the other schemes, we noted markedly faster DBE dissociation in the *Cavity* scheme (Figure S30), in line with the overrepresentation of bound states in the ASMDs. Interestingly, we found that the computed *k*_*d*_ values from all schemes were in good agreement with the experimentally determined one (Figure 4C). However, the computed values from the *Cavity* scheme were not well converged. The obtained less than an order of magnitude differences between simulations and experiments are still not common even in the case of much less complex biomolecular systems^64,65^.

### 3.3. Exploration of different transport paths by substrate

Next, we investigated the utilization of individual transport pathways of LinB86 by the substrate DBE. Initially, we attempted to match the substrate migration traces to the tunnel ensembles using Transport Tools library^67^. However, we could observe only very few complete migration events of DBE molecule between the bulk solvent and the active site of LinB86 (Table S3), with replicate 2 of the *Cavity* scheme capturing 33 transport events of DBE via known tunnels. Since such data cannot provide sufficient inference, we have followed by considering a simplified transition of DBE molecule through the tunnel bottleneck only, which corresponds to the least favorable region along the migration path and hence controls the transport rates^9,60,66^. Considering the distances of DBE to the COM of bottleneck residues of each tunnel and the bottom of the active site cavity (Figure S16), we have traced the location of DBE in all simulations, focusing on the frames where DBE came close to any of the bottlenecks and whether if passed through them.

A thorough investigation of the transport in all schemes via particular tunnels revealed the following observations. We observed the highest total number of transitions for the *Tunnels* scheme, followed by *Cavity, Cavity&Bulk*, and finally, the lowest in the *Bulk* only (Table S4). The overall proportion of the particular tunnels being utilized to the total number of transitions was consistent across all schemes. The most frequently used tunnel was *p2*, followed by *p1b, p1a, p3* and finally mixed and unknown representing the smallest fraction of the data (Figure 5A). Interestingly, besides this trend consistency, schemes *Cavity&Bulk* and *Tunnels* displayed a higher percentage of *p2* tunnel as compared to the remaining schemes (Figure 5A), suggesting that the more complex seeding schemes enable relatively more efficient exploration of the longest and most complex branches of *p2* tunnel. In contrast, simplified schemes (*Bulk* and *Cavity*) tend to promote the sampling of more accessible primary conduits, *p1* tunnels, in agreement with their preferential utilization observed in TransportTools analyses of replicate 2 of the *Cavity* scheme (Table S3).

**Figure 5.**
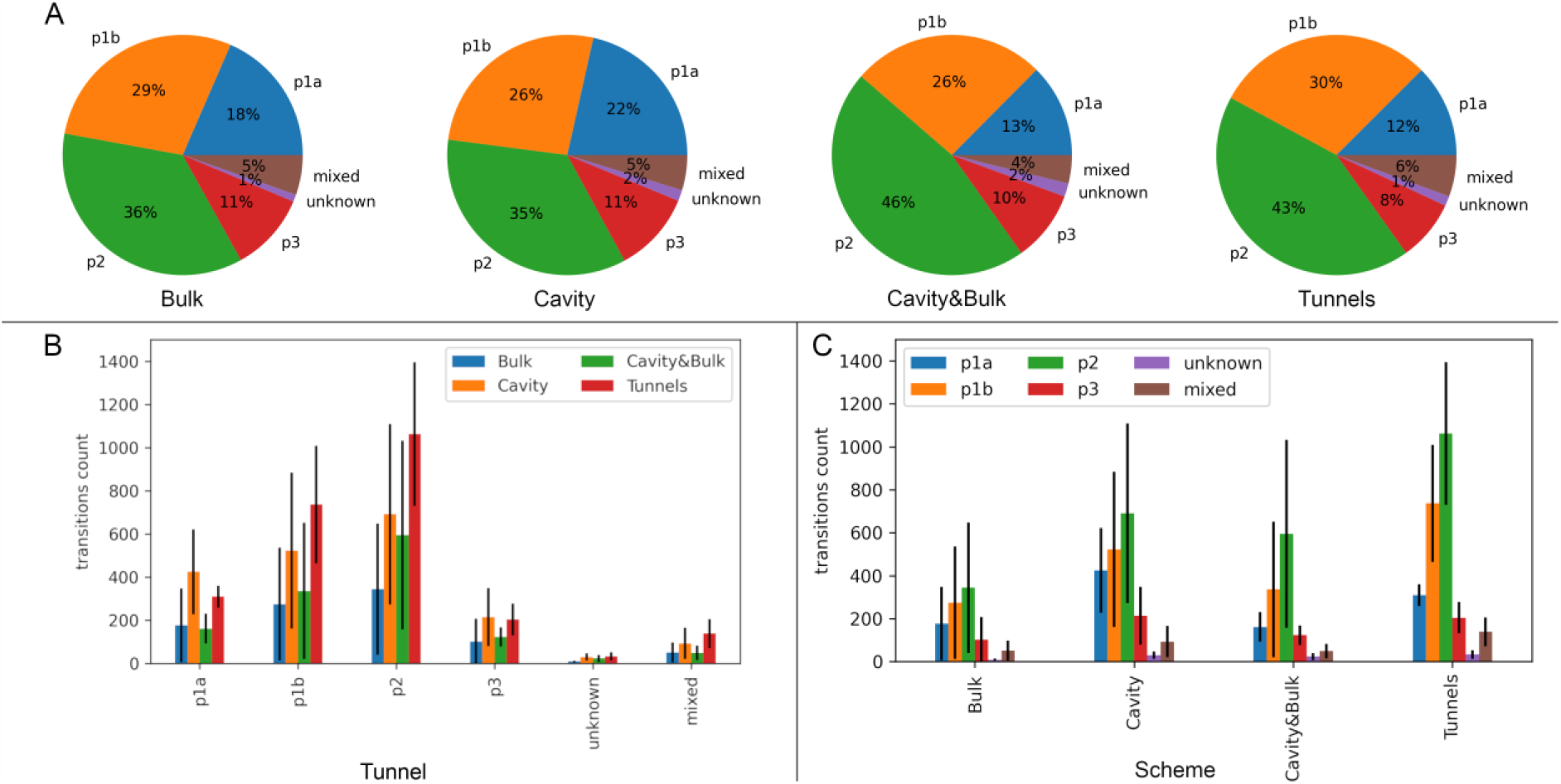
A statistical representation of tunnel utilization by the substrate for investigated seeding schemes. A) Relative utilization of particular tunnels for each scheme. B) Per tunnel average tunnel utilization. C) Per scheme average tunnel utilization. The data represents mean±stdev from the three replicates.

Besides this difference, the increased amount of transitions for particular schemes mainly came from the proportionally boosted sampling in each tunnel (Figure 5B). Importantly, considering the standard deviations calculated for combined statistics from three replicates for each scheme, it is clear that the *Tunnels* scheme presents the highest consistency from all tested schemes for all tunnels. This can be noticed when the transitions are considered for each run separately (Table S4). While the total sum of transitions for the *Bulk* scheme differs noticeably for particular replicates (1820, 49 and 1006 transitions for replica1, replica2 and replica3, respectively), the *Tunnels* scheme presents the lowest deviation when the three replicates are considered separately (replica1 – 2809, replica2 – 1680, replica3 – 2968). *Cavity* and *Cavity&Bulk* schemes fall between these two extremes and present similar consistency to each other (Figure 5C).

## 4. Conclusions

This study aimed to test the effect of different seeding schemes on effective sampling in MSM-driven ASMD simulations, providing meaningful insights into kinetic rates and mechanisms of the transport of the substrate DBE in LinB86 from its deeply buried active site to the solvent environment via multiple transport tunnels. The four designed seeding schemes allowed the positioning of DBE by focusing on applying more knowledge to tackle the sampling of regions with higher energy barriers. The ensuing ASMD simulations could construct the kinetic models with different levels of detail based on the employed seeding scheme. All simulations could explore the entire transport process, visiting unbound and bound states except for the *Bulk* scheme that could not reach the bound state in two replicates of 45 μs ASMDs. Conversely, the *Tunnels* scheme was most consistent in sampling different metastable states of substrate in the transport-relevant regions. The application of more information-rich *Tunnels* and *Cavity&Bulk* schemes led to the enhanced exploration of auxiliary *p2* and *p3* tunnels. In contrast, primary *p1* tunnels were preferred in ASMDs initiated from the other two schemes. *Tunnels* and *Cavity&Bulk* schemes also provided the most converged *k*_*d*_ values from the rates of DBE association and dissociation, sufficiently close to the experimental measurements despite the complexity of the kinetic model. We expect that analogous methodology can also be beneficial for defining effective collective variables enhanced sampling methods like metadynamics,^21,22^ and umbrella sampling^68^. Overall, the infusion of more knowledge into the initial seeds of ASMD simulations could render computational analyses of transport mechanisms in enzymes more consistent even for very complex biomolecular systems, having a clear potential to translate into faster rational protein design and drug development efforts.

## Author contributions

Conceptualization: J.B.; Data curation: D.K.S, Formal analysis: D.K.S., B.S. J.B.; Funding acquisition: J.B., Investigation: D.K.S. (Simulations and MSMs), B.S. (Transitions and ULS clustering); Methodology:

D.K.S. (Simulations and MSMs), B.S. (Transitions and ULS clustering); Project administration: J.B.; Resources: J.B.; Software: J.B.; Supervision: J.B.; Validation: J.B.; Writing – original draft: D.K.S. (Simulations and MSMs), B.S. (Transitions and ULS clustering); Writing – review & editing: J.B.

## Acknowledgments

This work was supported by the National Science Centre, Poland (grant no. 2017/26/E/NZ1/00548). D.K.S. and B.S. were scholarship recipients provided by POWER projects POWR.03.02.00-00-I006/17 and POWR.03.02.00-00-I022/16, respectively. The simulations were performed in the Poznan supercomputing center.

