## Supporting information for "Incorporating prior knowledge to seeds of adaptive sampling molecular dynamics simulations of ligand transport in enzymes with buried active sites"

**Table S1. Properties of tunnel identified in LinB86 simulation using CAVER.**

| *ID* | *No* | *No_snaps* | *Avg_BR* | *Avg_BR SD* | *Max_BR* | *Avg_L* | *Avg_L SD* | *Avg_C* | *Avg_C SD* | *Priority* | *Avg_throughput* | *Avg_throughput SD* |
|---|---|---|---|---|---|---|---|---|---|---|---|---|
| ***p1a*** | 121 | 121 | 1.07 | 0.153 | 1.61 | 16.01 | 2.542 | 1.403 | 0.167 | 0.11104 | 0.45885 | 0.09486 |
| ***p1b*** | 146 | 146 | 1.251 | 0.233 | 1.75 | 13.608 | 1.723 | 1.277 | 0.138 | 0.17275 | 0.5916 | 0.10193 |
| ***p2a*** | 231 | 231 | 1.191 | 0.201 | 1.82 | 19.985 | 2.535 | 1.271 | 0.108 | 0.21728 | 0.4703 | 0.09811 |
| ***p2c*** | 237 | 237 | 1.045 | 0.107 | 1.58 | 22.873 | 2.676 | 1.406 | 0.156 | 0.17568 | 0.37064 | 0.06156 |
| ***p2b*** | 184 | 184 | 1.093 | 0.17 | 1.61 | 17.145 | 2.413 | 1.409 | 0.159 | 0.17297 | 0.47004 | 0.0844 |
| ***p2d*** | 164 | 164 | 1.044 | 0.111 | 1.49 | 25.6 | 3.137 | 1.754 | 0.285 | 0.10949 | 0.33382 | 0.0682 |
| ***p3*** | 97 | 97 | 1.019 | 0.11 | 1.38 | 16.284 | 1.955 | 1.255 | 0.103 | 0.08738 | 0.45042 | 0.06983 |

**ID: tunnel cluster ranked based on priority, No: Total number of tunnels, No_snaps: Number of snapshots, Avg_BR: Average bottleneck radius (Å), Max_BR: Maximum bottleneck radius (Å), Avg_L: Average tunnel length (Å), Avg_C: Average tunnel curvature, Priority: Tunnel priority calculated by averaging tunnel throughputs over all snapshots, Avg_throughput: Average tunnel throughput.*

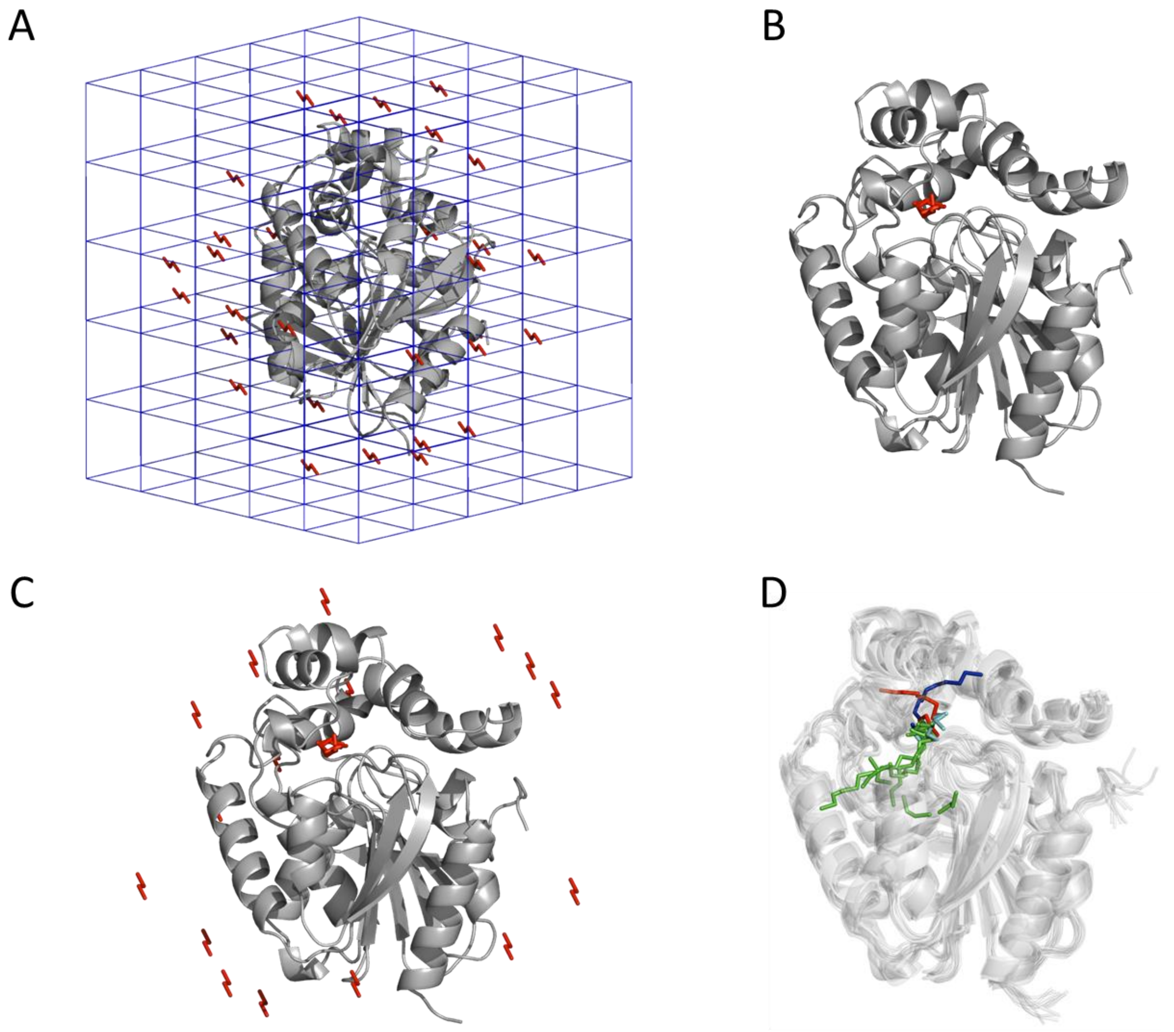


**Figure S1. Initial seeds of the DBE molecule around the LinB86 enzyme used in four investigated schemes.** A) *Bulk*, B) *Cavity*, C) *Cavity&Bulk,* and D) *Tunnels* schemes. In the *Tunnels* scheme, not only positions of DBE molecules were seeded, but also the corresponding structure of LinB86 harboring appropriate tunnel fragments was used. Protein structure is shown as gray (static) or white (ensemble) cartoon. Positions of DBE are shown in red sticks, except for the *Tunnels* scheme in which sticks are colored according to the tunnel seeded: *p1a* (blue), *p1b* (cyan), *p2* (green), and *p3* (red).

**Text S1 Protocol for MD simulation for tunnel calculations used during seeding with scheme *Tunnels:*** For MD setup, AMBER18[1] package was used and the crystal structure of LinB86 was retrieved from the PDB database (ID 5LKA). The periodic boundary conditions were maintained by particle mesh Ewald method[2] with the SHAKE algorithm for 4 fs time-step and hydrogen mass repartitioning method[3]. For minimization, the system was subjected to five rounds of minimization consisting of 500 steps of steepest descent minimization. After minimization, the system was gradually heated at canonical NVT ensemble from 0 K to 200 K with restrained heavy protein atoms for 20 ps using the Langevin thermostat. Then the system was equilibrated for 2 ns in three rounds. First using harmonic restraints, the system was gradually heated to 310 K within 100 ps and keeping constant temperature for another 900 ps in canonical NVT ensemble. Secondly, the system was subjected to constant pressure and temperature (NPT) while restraining the backbone atoms for 1 ns, using a weak-coupling barostat. Finally, 100 ns of NPT simulation without restraints was performed.

The MD trajectory was analyzed to identify geometric tunnels by CAVER 3.0.1[4,5] with the starting points of tunnels defined by the following the active site residues Asn38, Asp108, Trp109, and His272, using a probe radius of 0.9 Å, shell radius 3 Å and shell depth 4 Å. The obtained tunnels were clustered by hierarchical clustering with a clustering threshold of 3.5 Å. For each known tunnel, i.e., *p1a*, *p1b*, *p2,* and *p3* (Table S1), the 100 most opened tunnels were selected for positioning the DBE molecule along the entire tunnel length by the CaverDock program[6]. As a result, sets of DBE positions along the tunnel and the corresponding upper-bound interaction energies were generated for each tunnel ensemble. These energies were converted to approximate maximal migration barriers for tunnel segments of 1 Å length by finding the highest energy in the segment and subtracting the global minimal interaction energy found among the entire tunnel ensemble (Figure S2-S8).

Next, we have generated composite tunnels with putatively minimal energy costs for DBE migration through the respective tunnel conformations (Figure S9). For each tunnel segment, the tunnel conformation with the lowest migration barriers for DBE transport was selected unless the barrier of the previous optimal tunnel conformation was within 1 kcal/mol for this segment too. In such a case, the same tunnel conformation was retained to avoid frequent switching between the tunnel conformations. This procedure resulted in the generation of a subset of tunnel segments with the corresponding protein conformations with favorably bound DBE along the whole tunnel length (Figure S1D).

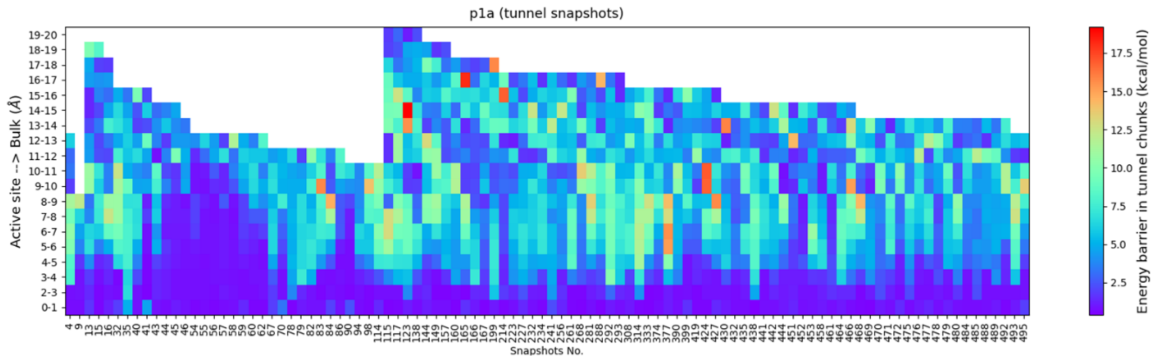


**Figure S2. DBE migration energy barriers profile of *p1a* tunnel ensemble.** The values correspond to the maximal interaction energy in each bin (1 Å length) predicted by CaverDock decreased by the global minimal interaction energy in the tunnel ensemble (-3.1 kcal/mol).

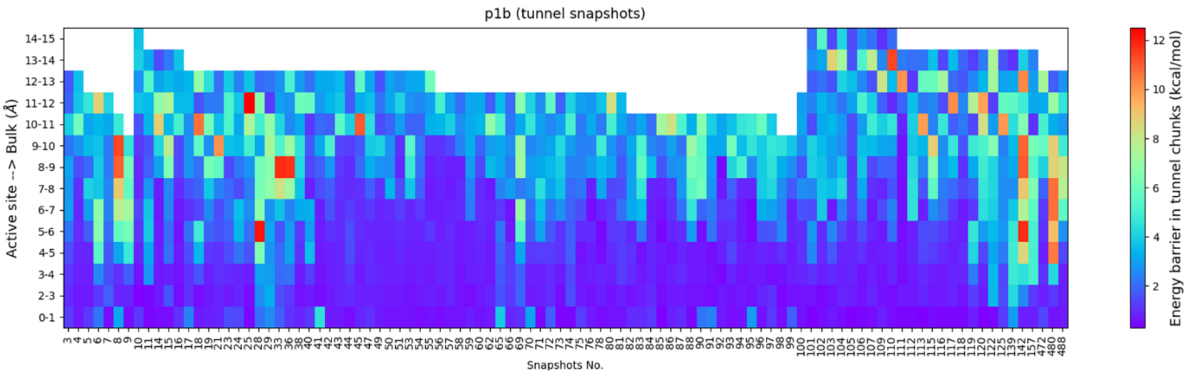


**Figure S3. DBE migration energy barriers profile of *p1b* tunnel ensemble.** The values correspond to the maximal interaction energy in each bin (1 Å length) predicted by CaverDock decreased by the global minimal interaction energy in the tunnel ensemble (-3.0 kcal/mol).

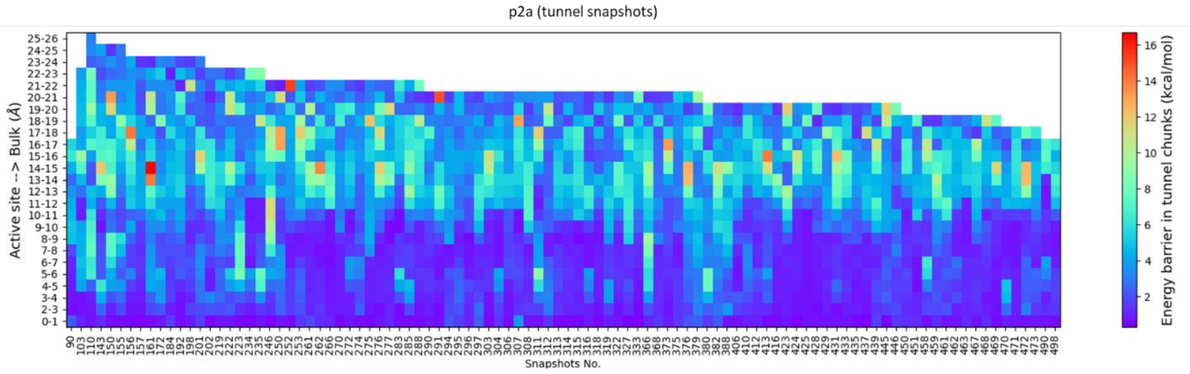


**Figure S4. DBE migration energy barriers profile of *p2a* tunnel ensemble.** The values correspond to the maximal interaction energy in each bin (1 Å length) predicted by CaverDock decreased by the global minimal interaction energy in the tunnel ensemble (-3.0 kcal/mol).

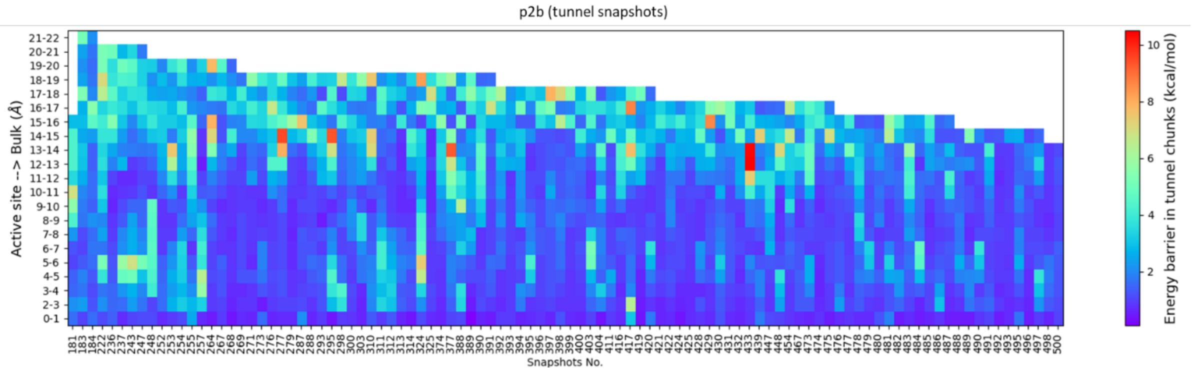


**Figure S5. DBE migration energy barriers profile of *p2b* tunnel ensemble.** The values correspond to the maximal interaction energy in each bin (1 Å length) predicted by CaverDock decreased by the global minimal interaction energy in the tunnel ensemble (-3.0 kcal/mol).

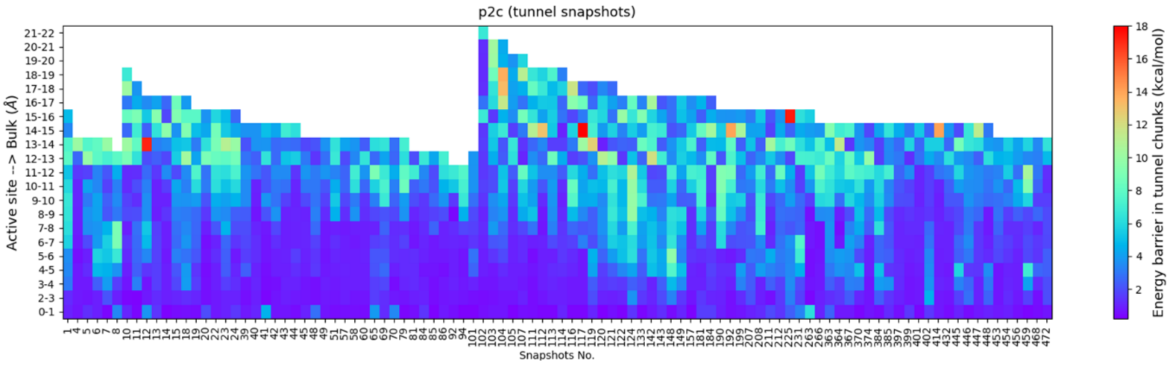


**Figure S6. DBE migration energy barriers profile of *p2c* tunnel ensemble.** The values correspond to the maximal interaction energy in each bin (1 Å length) predicted by CaverDock decreased by the global minimal interaction energy in the tunnel ensemble (-3.0 kcal/mol).

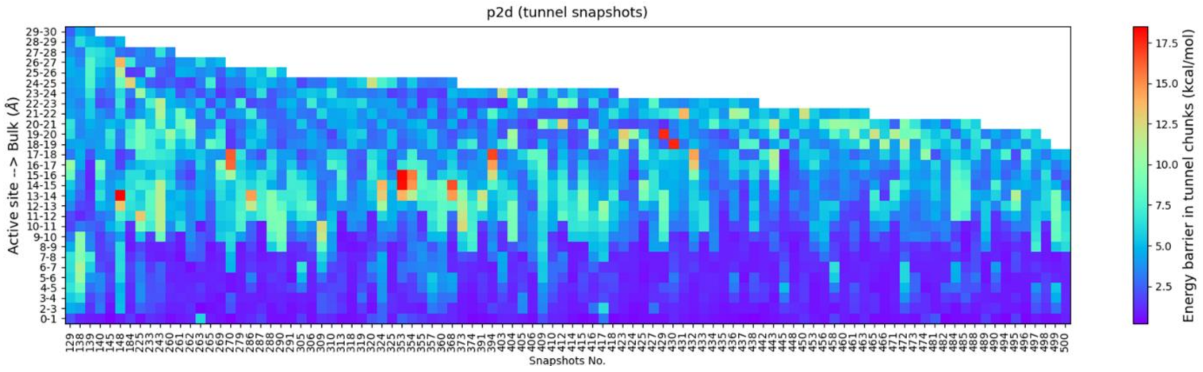


**Figure S7. DBE migration energy barriers profile of *p2d* tunnel ensemble.** The values correspond to the maximal interaction energy in each bin (1 Å length) predicted by CaverDock decreased by the global minimal interaction energy in the tunnel ensemble (-3.0 kcal/mol).

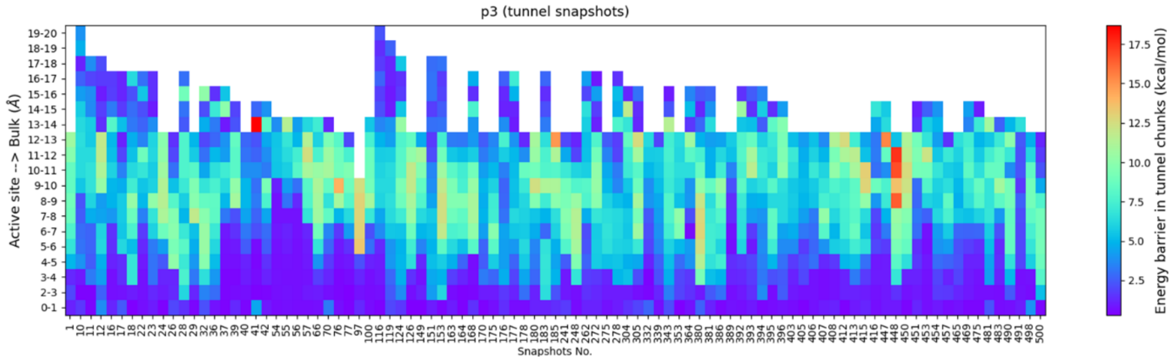


**Figure S8. DBE migration energy barriers profile of *p3* tunnel ensemble.** The values correspond to the maximal interaction energy in each bin (1 Å length) predicted by CaverDock decreased by the global minimal interaction energy in the tunnel ensemble (-3.0 kcal/mol).

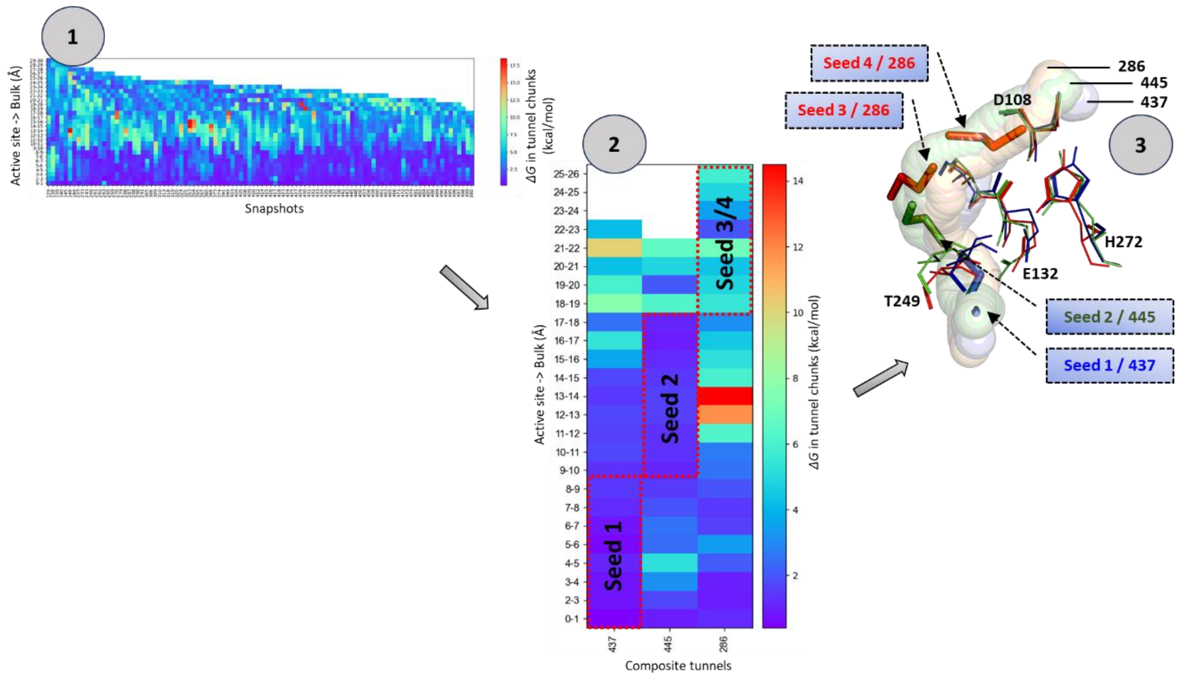


**Figure S9. Creation of composite tunnel with corresponding bound poses of DBE molecule for seeding**. *Step 1:* Compiling the ensemble of tunnels and maximal migration barriers for DBE molecule computed based on CaverDock energy profiles for each tunnel. *Step 2:* generation of composite tunnels (from several favorable segments of tunnels from the ensemble). Finally, in *Step 3*: the positions of ligands are selected for seeding based on the composite tunnel profiles by placing the DBE in favorable positions.

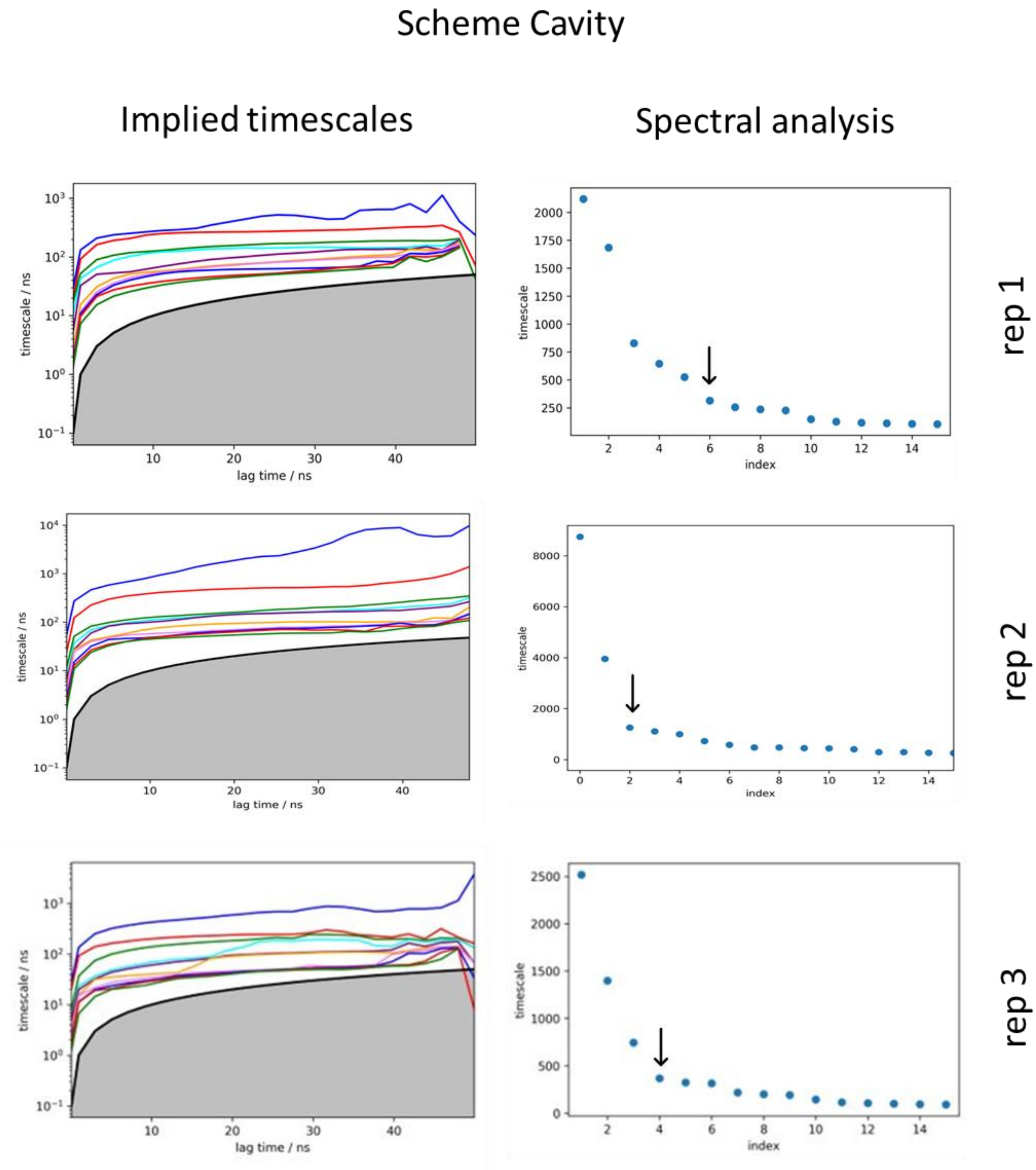


**Figure S10. Implied time scales of MSM generated for three replicates of the *Cavity* scheme and corresponding spectral separation analysis.** The arrow indicates the last point considered for deciding the number of metastable states.

#### Scheme Cavity&Bulk

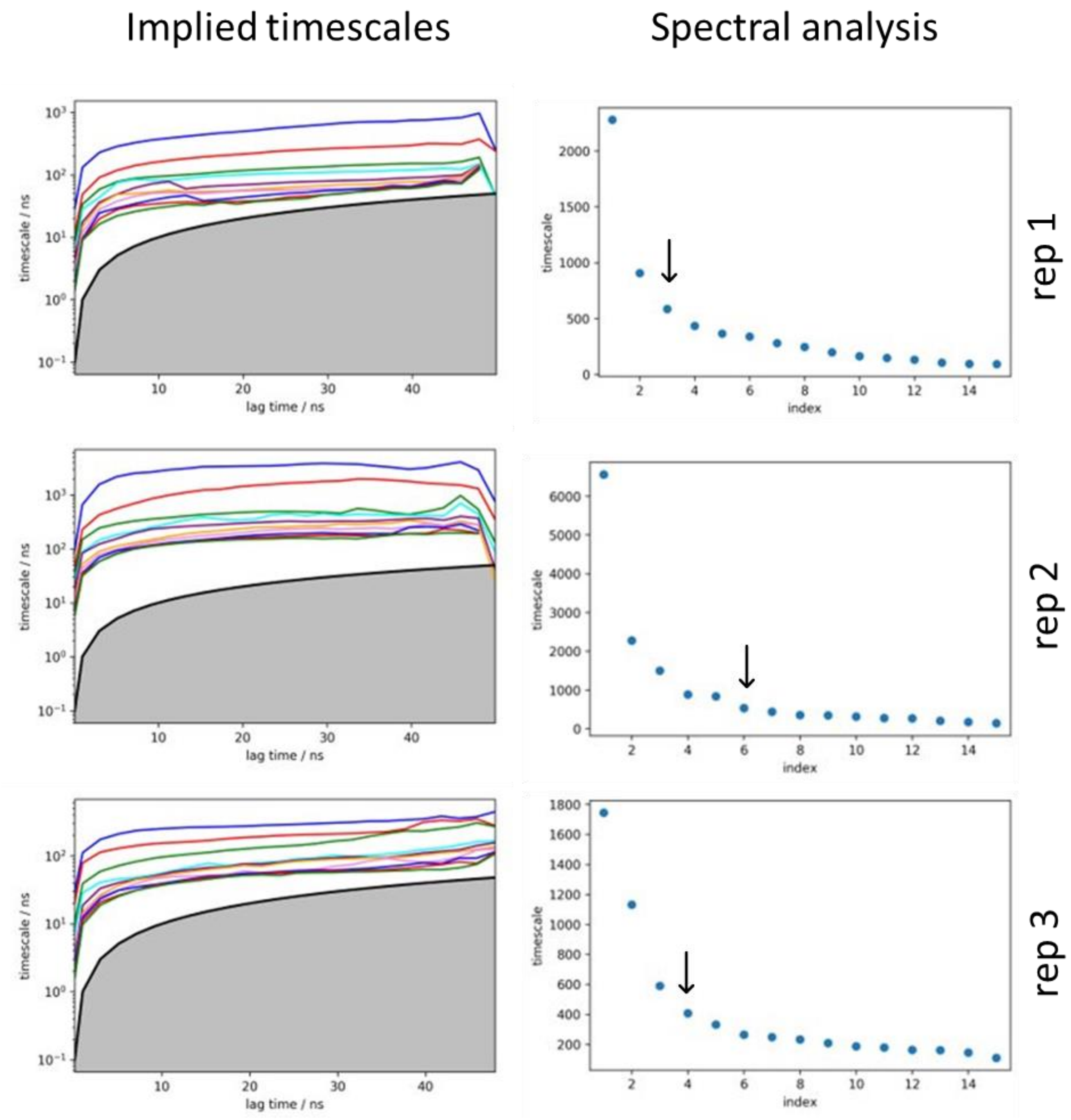


**Figure S11. Implied time scales of MSM generated for three replicates of the *Cavity&Bulk* scheme and corresponding spectral separation analysis.** The arrow indicates the last point considered for deciding the number of metastable states.

### Scheme Tunnels

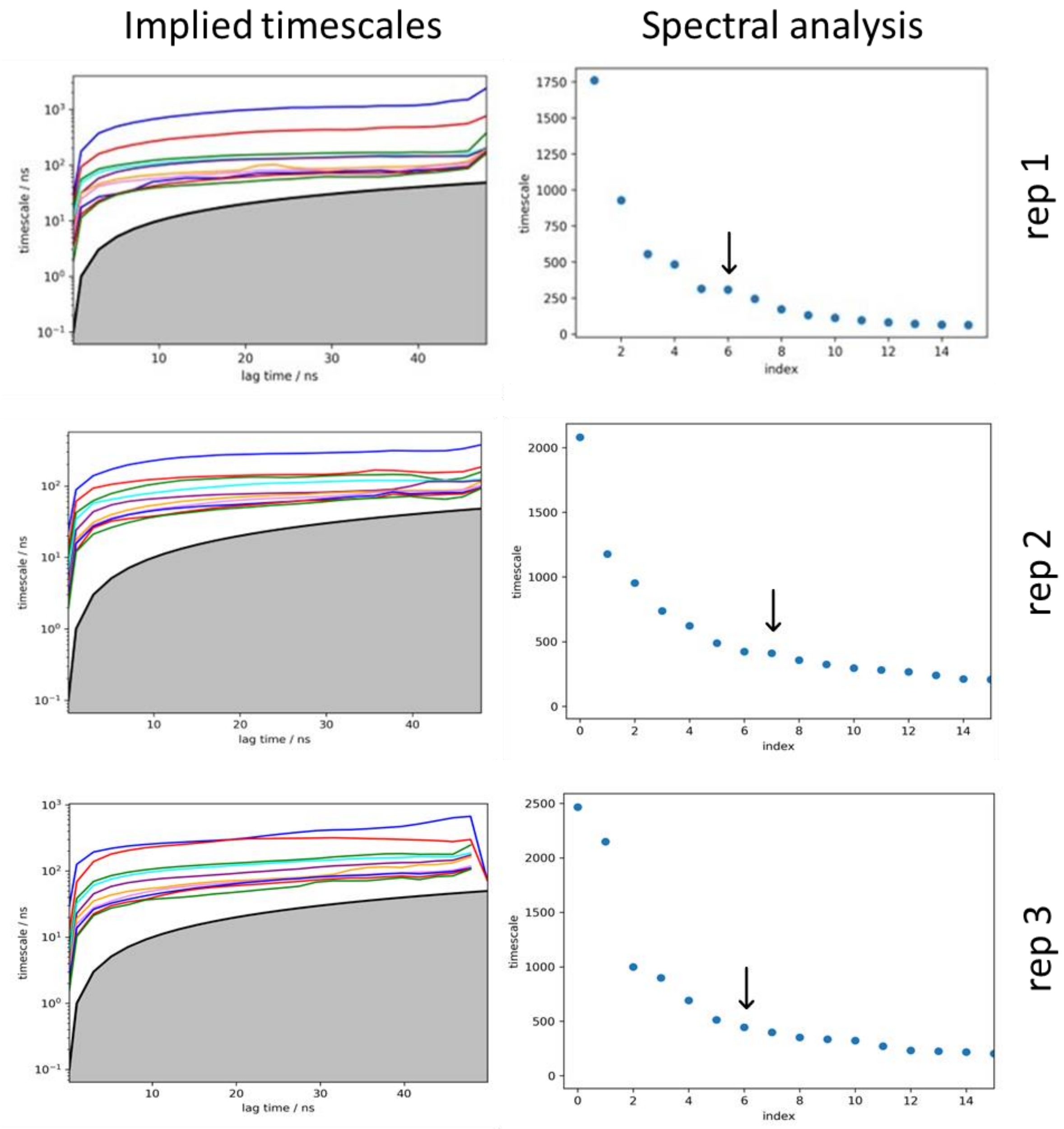


**Figure S12. Implied time scales of MSM generated for three replicates of the *Tunnels* scheme and corresponding spectral separation analysis.** The arrow indicates the last point considered for deciding the number of metastable states.

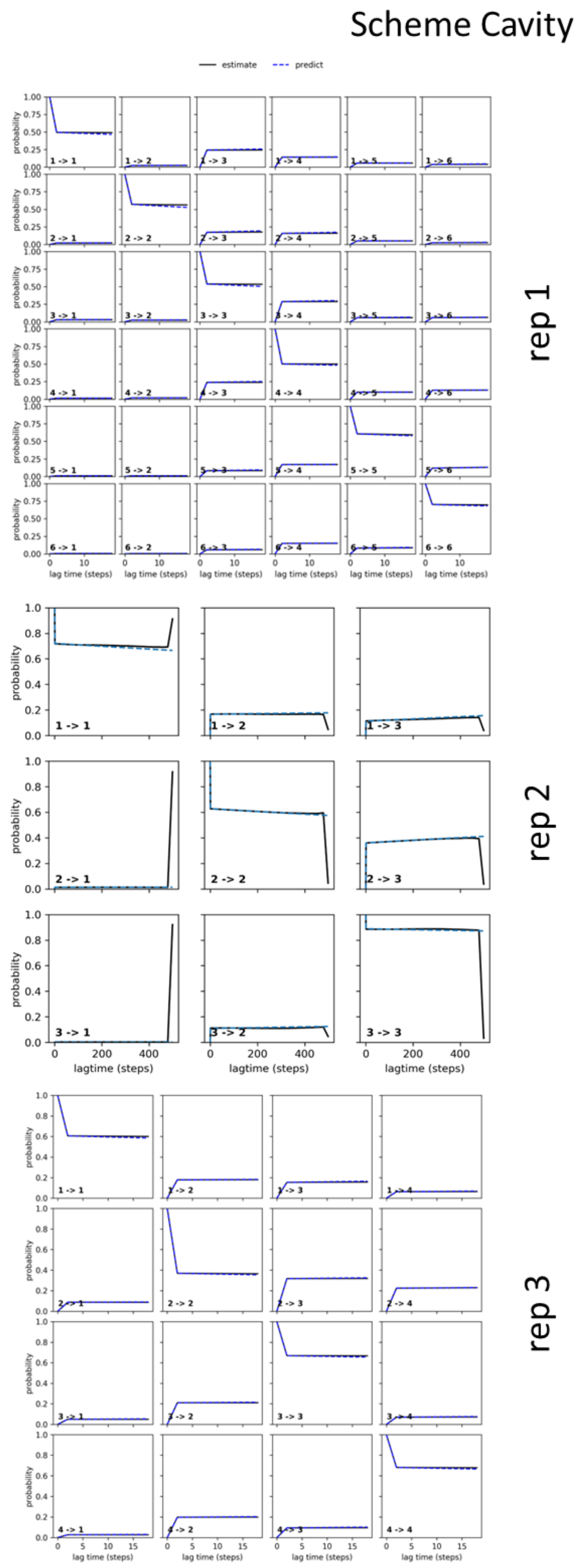


**Figure S13. Chapman-Kolmogorov tests for three replicates of the *Cavity* scheme.** Plots show that MSMs correctly represent original MD simulation data.

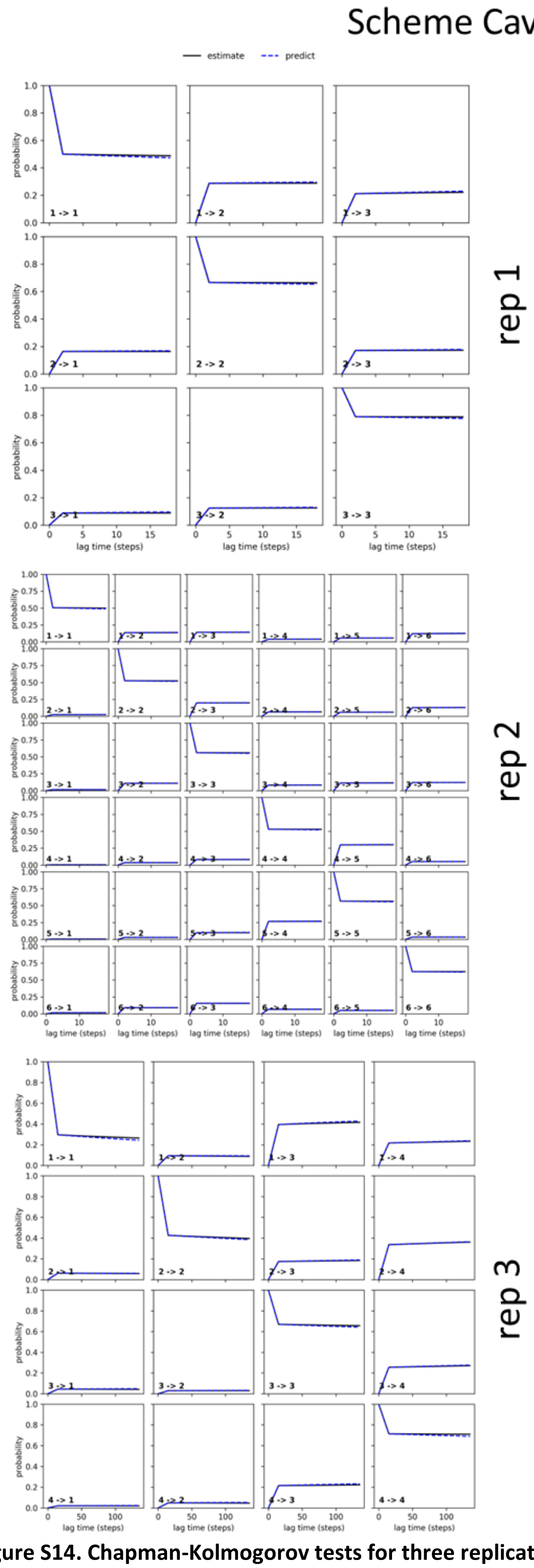


**Figure S14. Chapman-Kolmogorov tests for three replicates of the *Cavity&Bulk* scheme.** Plots show that MSMs correctly represent original MD simulation data.

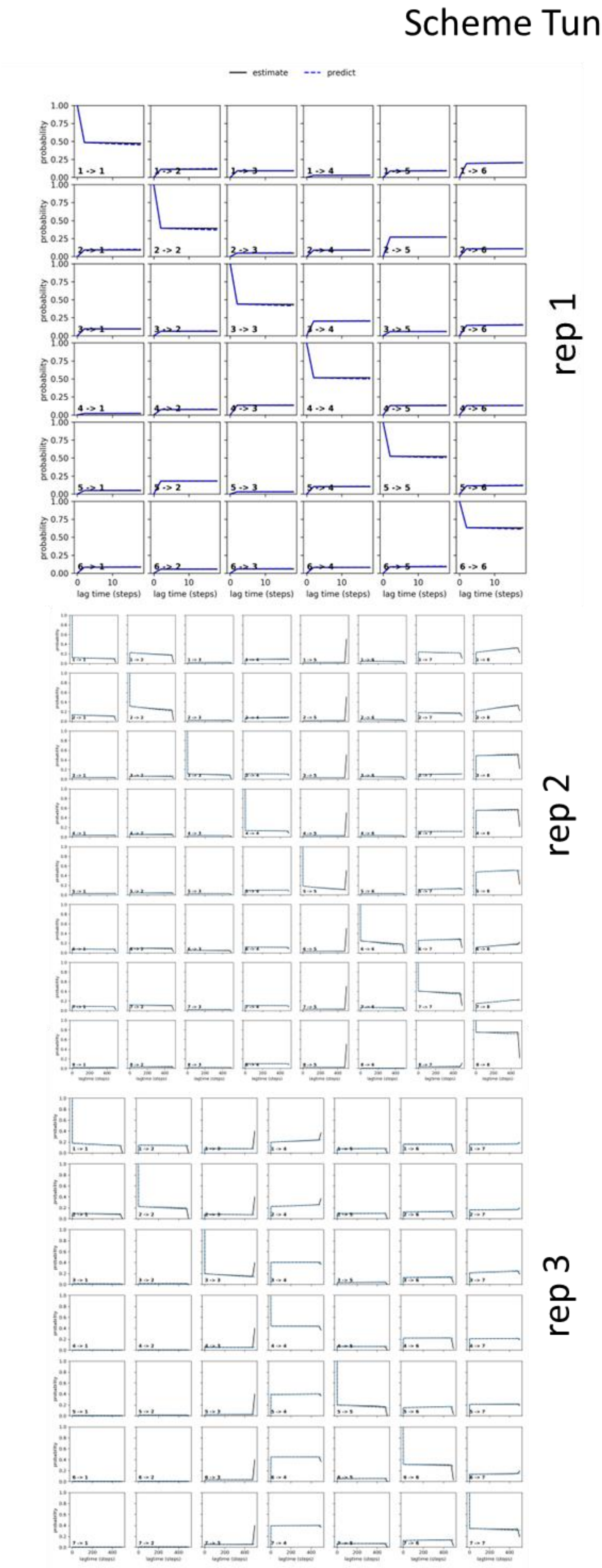


**Figure S15. Chapman-Kolmogorov tests for three replicates of the *Tunnels* scheme.** Plots show that MSMs correctly represent original MD simulation data.

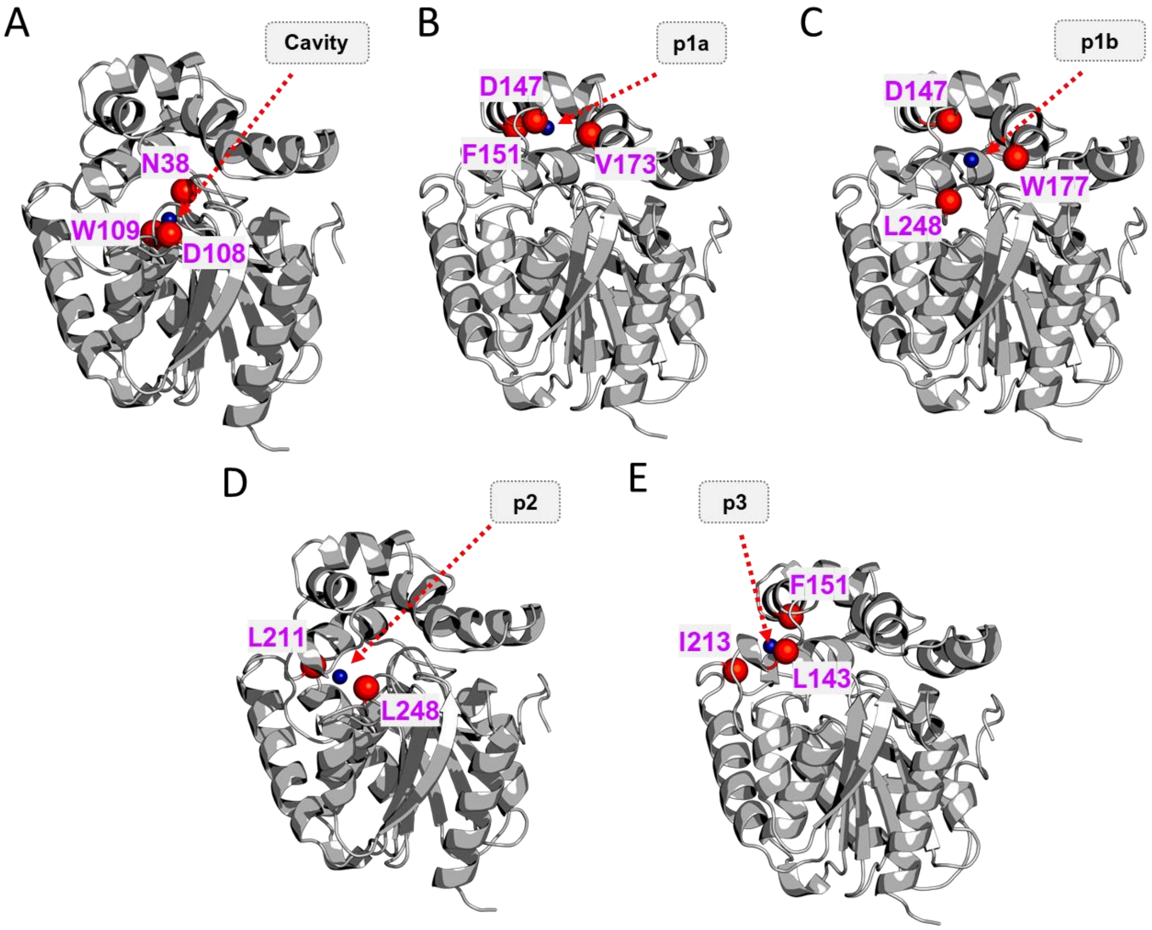


**Figure S16. The COM of residues used to characterize DBE locations within the LinB86 structure.** A) three catalytic residues defining the bottom of the active site cavity, B) residues defining the bottleneck of the *p1a* tunnel, C) residues defining the bottleneck of the *p1b* tunnel, D) residues defining the bottleneck of the *p2* tunnels, and E) residues defining the bottleneck of *p3* tunnel. The COMs are shown as blue spheres and the positions of the residues are defined by their CA atoms, shown as red spheres.

**Table S2. The number of ASMD simulations successfully completed per scheme and replicate.**

| Schemes | Replicates | Number of simulations | Total time [µs] |
|---|---|---|---|
| Bulk | 1 | 898 | 44.90 |
| | 2 | 900 | 45.00 |
| | 3 | 899 | 44.95 |
| Cavity | 1 | 888 | 44.40 |
| | 2 | 900 | 45.00 |
| | 3 | 897 | 44.85 |
| Cavity&Bulk | 1 | 899 | 44.95 |
| | 2 | 900 | 45.00 |
| | 3 | 891 | 44.55 |
| Tunnels | 1 | 886 | 44.30 |
| | 2 | 888 | 44.40 |
| | 3 | 897 | 44.85 |

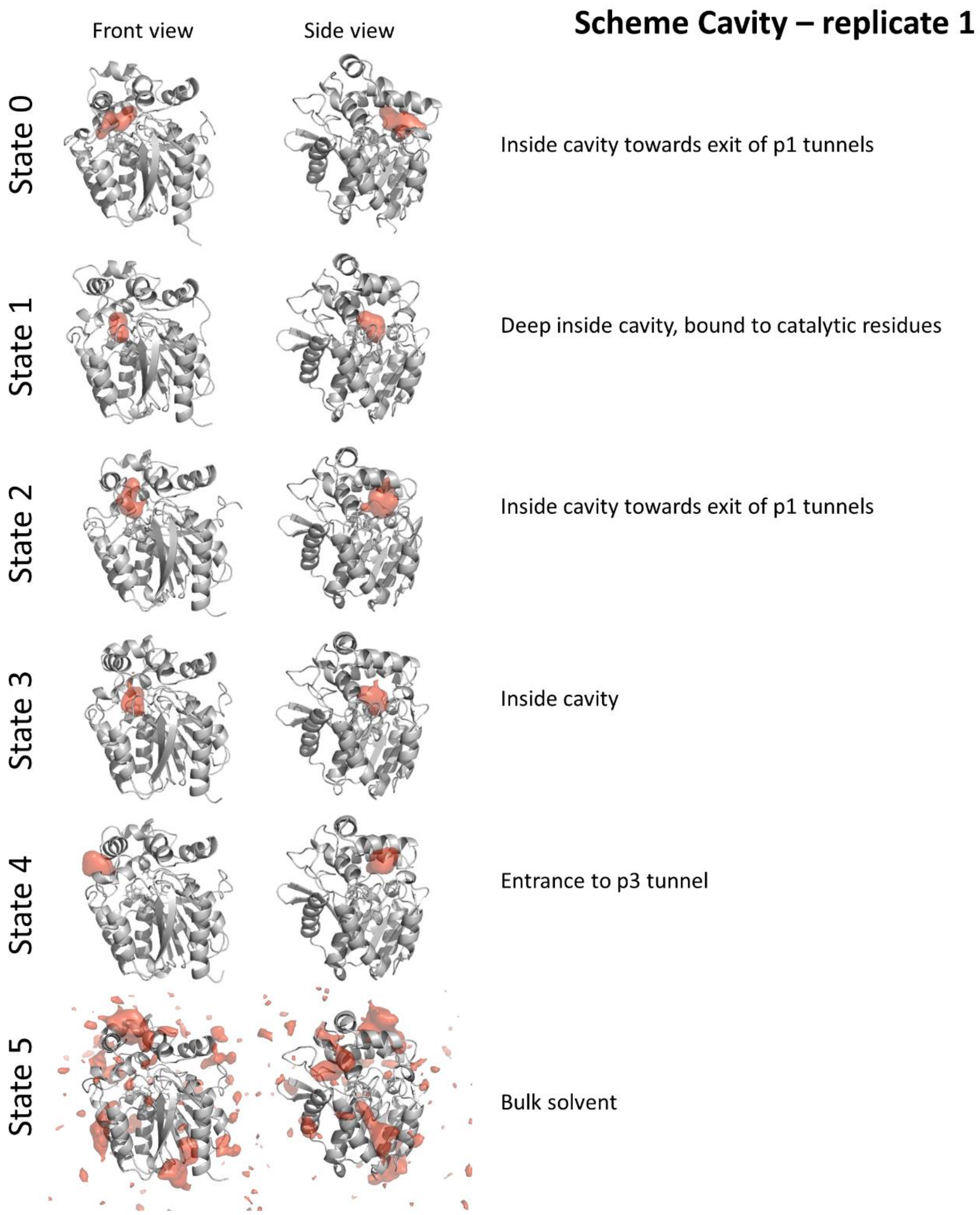


**Figure S17. Overview metastable states derived from MSM of *Cavity* scheme replicate 1.** Protein structure is shown as a gray cartoon while the region occupied by DBE molecule in 20 % (1 % for bulk solvent state) of 1000 structures representing given metastable state is shown as red surface.

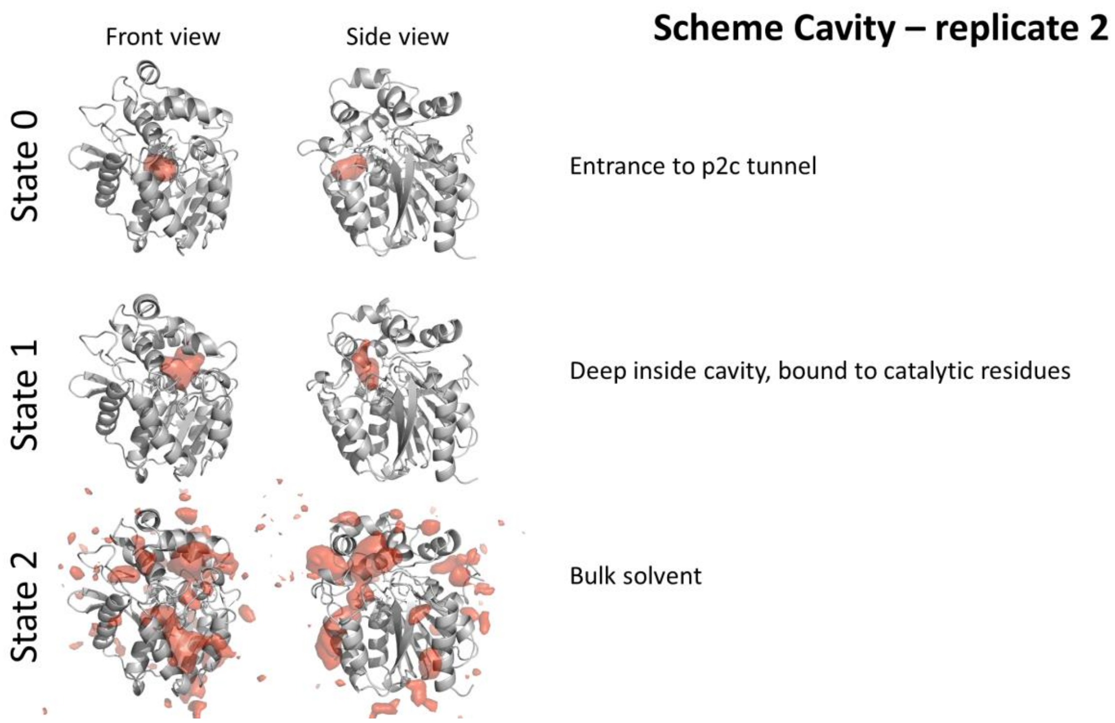


**Figure S18. Overview metastable states derived from MSM of *Cavity* scheme replicate 2.** Protein structure is shown as a gray cartoon while the region occupied by DBE molecule in 20 % (1 % for bulk solvent state) of 1000 structures representing given metastable state is shown as red surface.

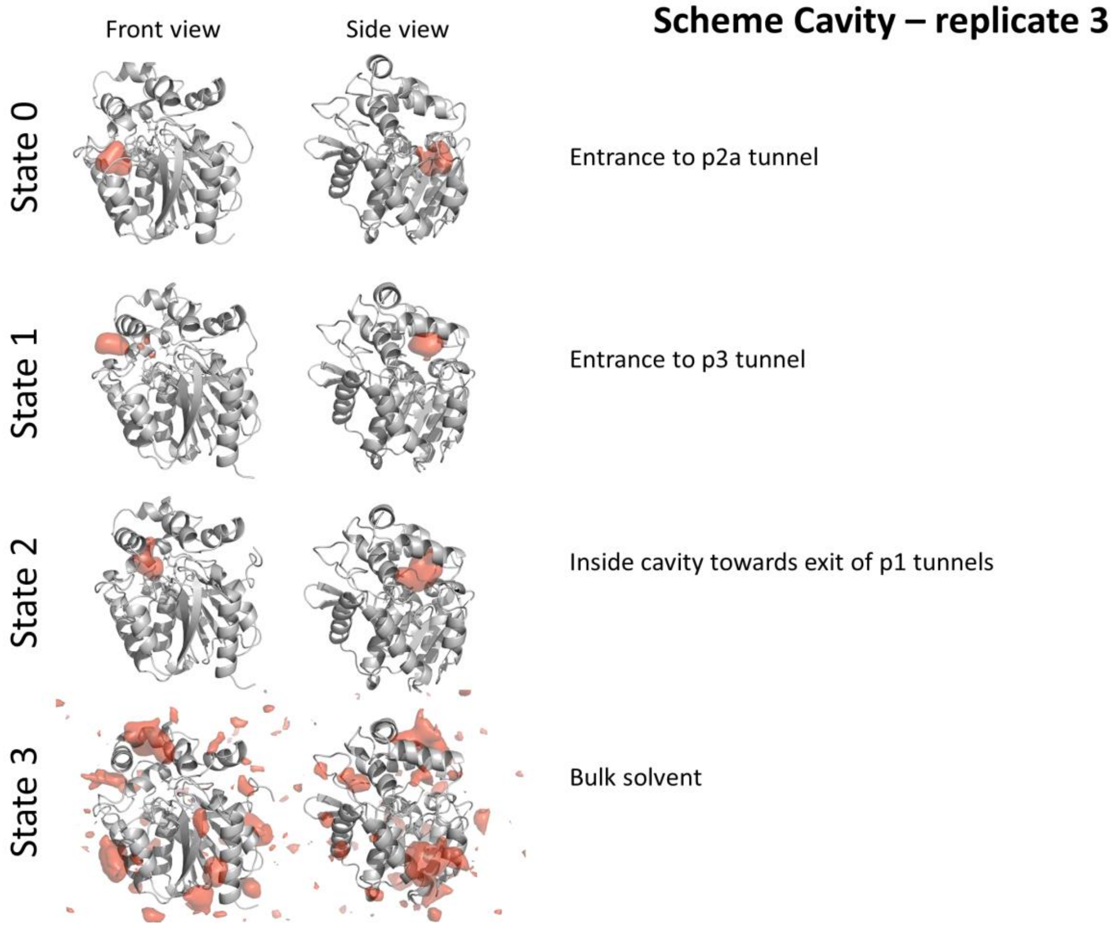


**Figure S19. Overview metastable states derived from MSM of *Cavity* scheme replicate 3.** Protein structure is shown as a gray cartoon while the region occupied by DBE molecule in 20 % (1 % for bulk solvent state) of 1000 structures representing given metastable state is shown as red surface.

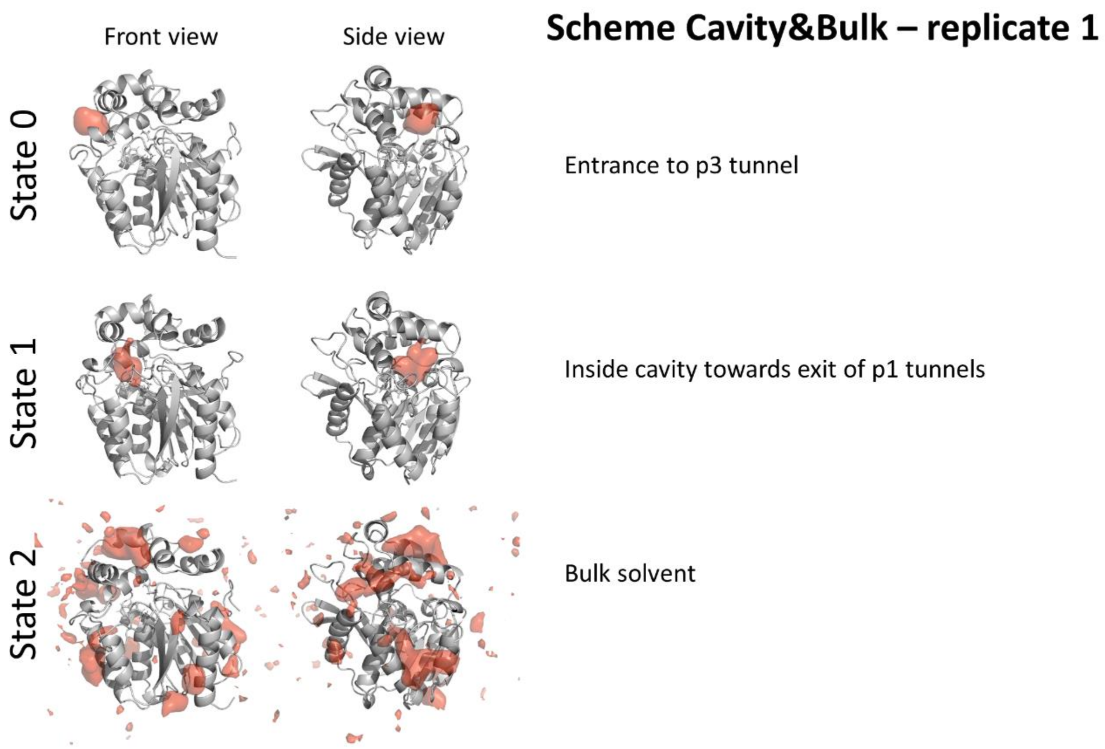


**Figure S20. Overview metastable states derived from MSM of *Cavity&Bulk* scheme replicate 1.** Protein structure is shown as a gray cartoon while the region occupied by DBE molecule in 20 % (1 % for bulk solvent state) of 1000 structures representing given metastable state is shown as red surface.

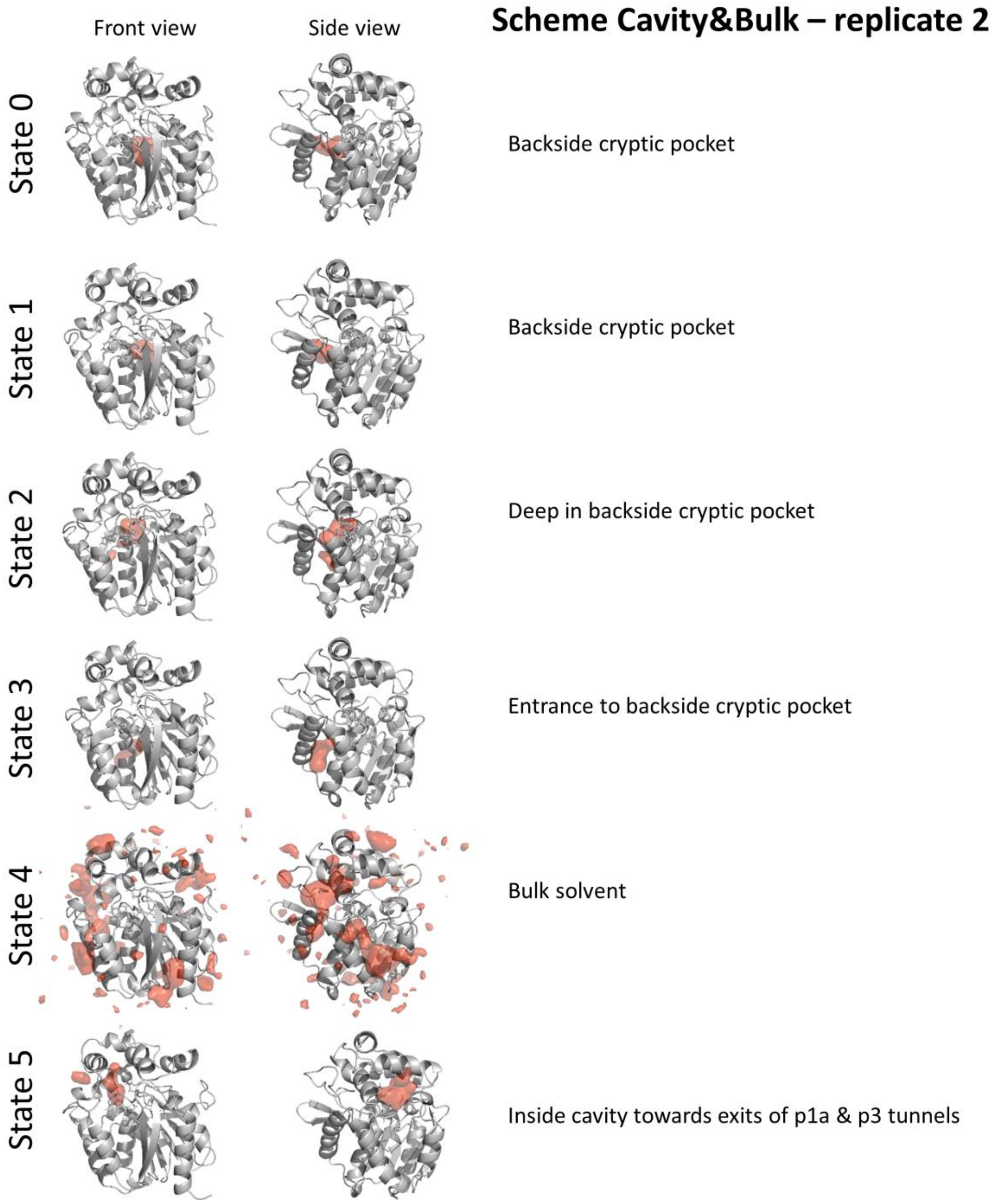


**Figure S21. Overview metastable states derived from MSM of *Cavity&Bulk* scheme replicate 2.** Protein structure is shown as a gray cartoon while the region occupied by DBE molecule in 20 % (1 % for bulk solvent state) of 1000 structures representing given metastable state is shown as red surface.

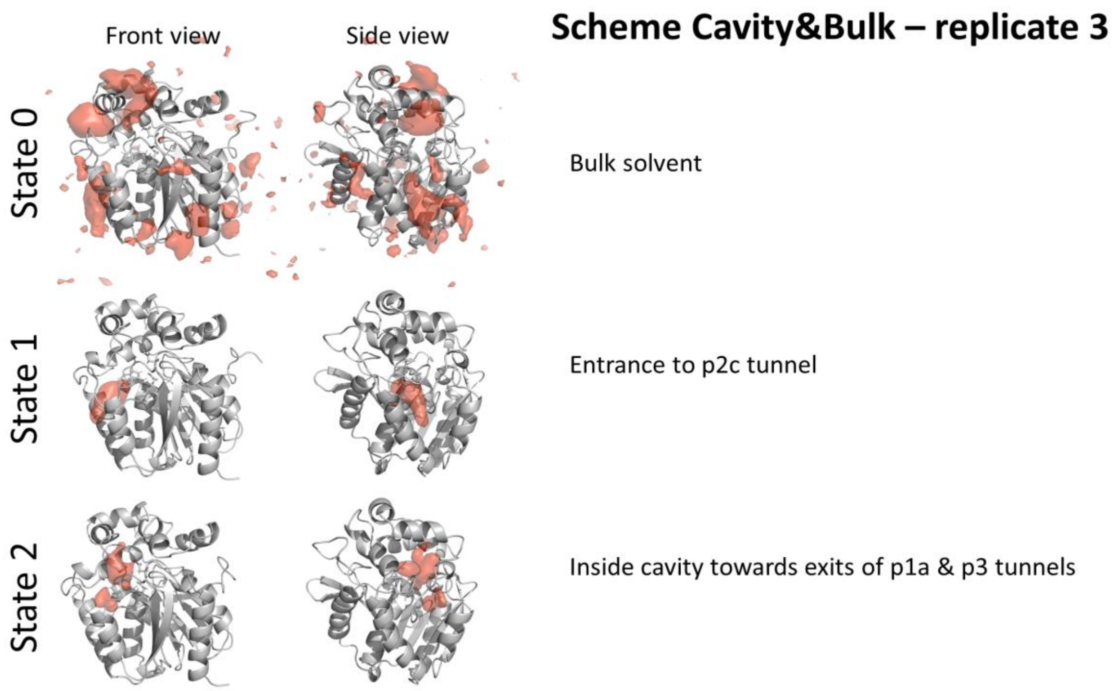


**Figure S22. Overview metastable states derived from MSM of *Cavity&Bulk* scheme replicate 3.** Protein structure is shown as a gray cartoon while the region occupied by DBE molecule in 20 % (1 % for bulk solvent state) of 1000 structures representing given metastable state is shown as red surface.

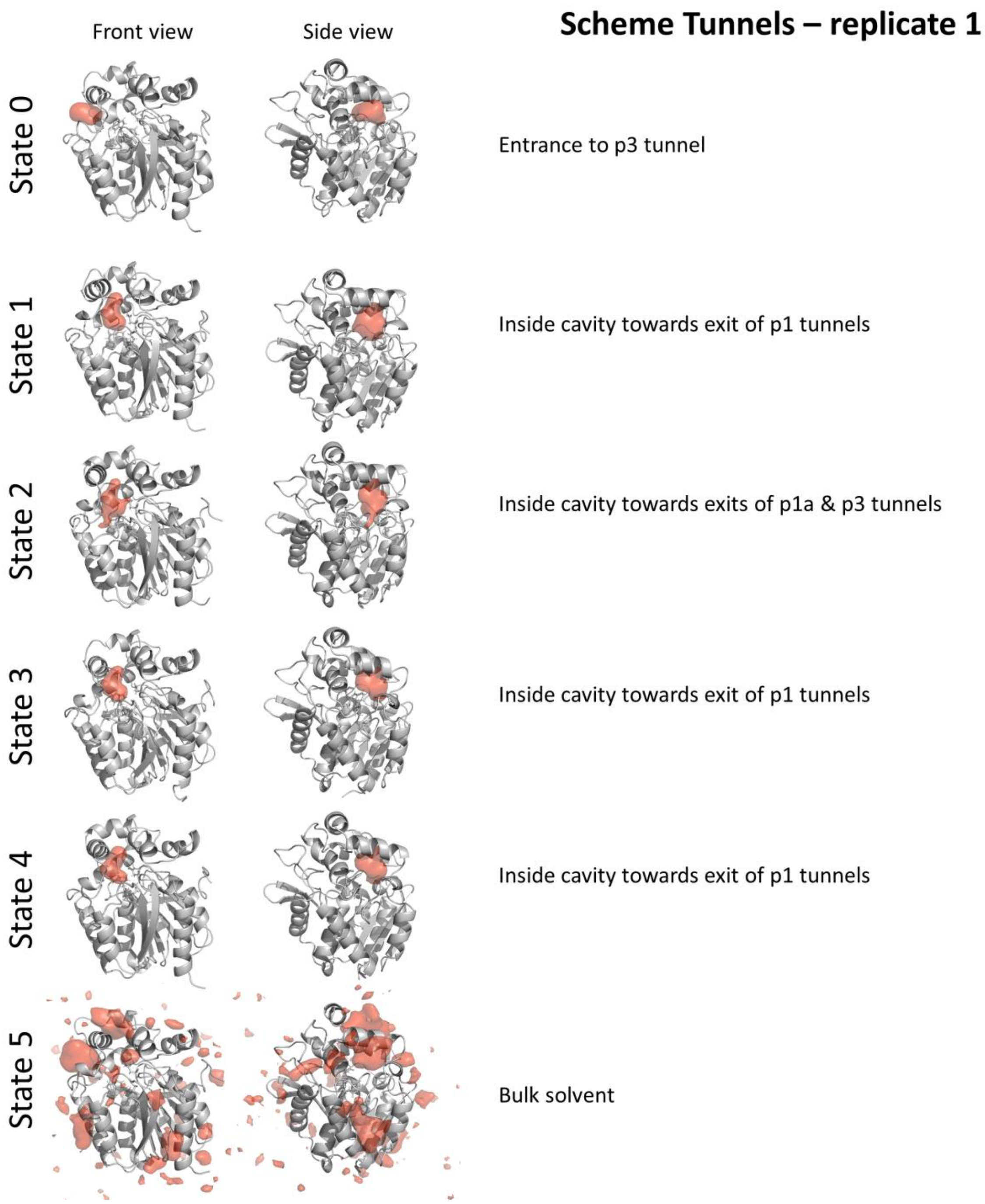


**Figure S23. Overview metastable states derived from MSM of *Tunnels* scheme replicate 1.** Protein structure is shown as a gray cartoon while the region occupied by DBE molecule in 20 % (1 % for bulk solvent state) of 1000 structures representing given metastable state is shown as red surface.

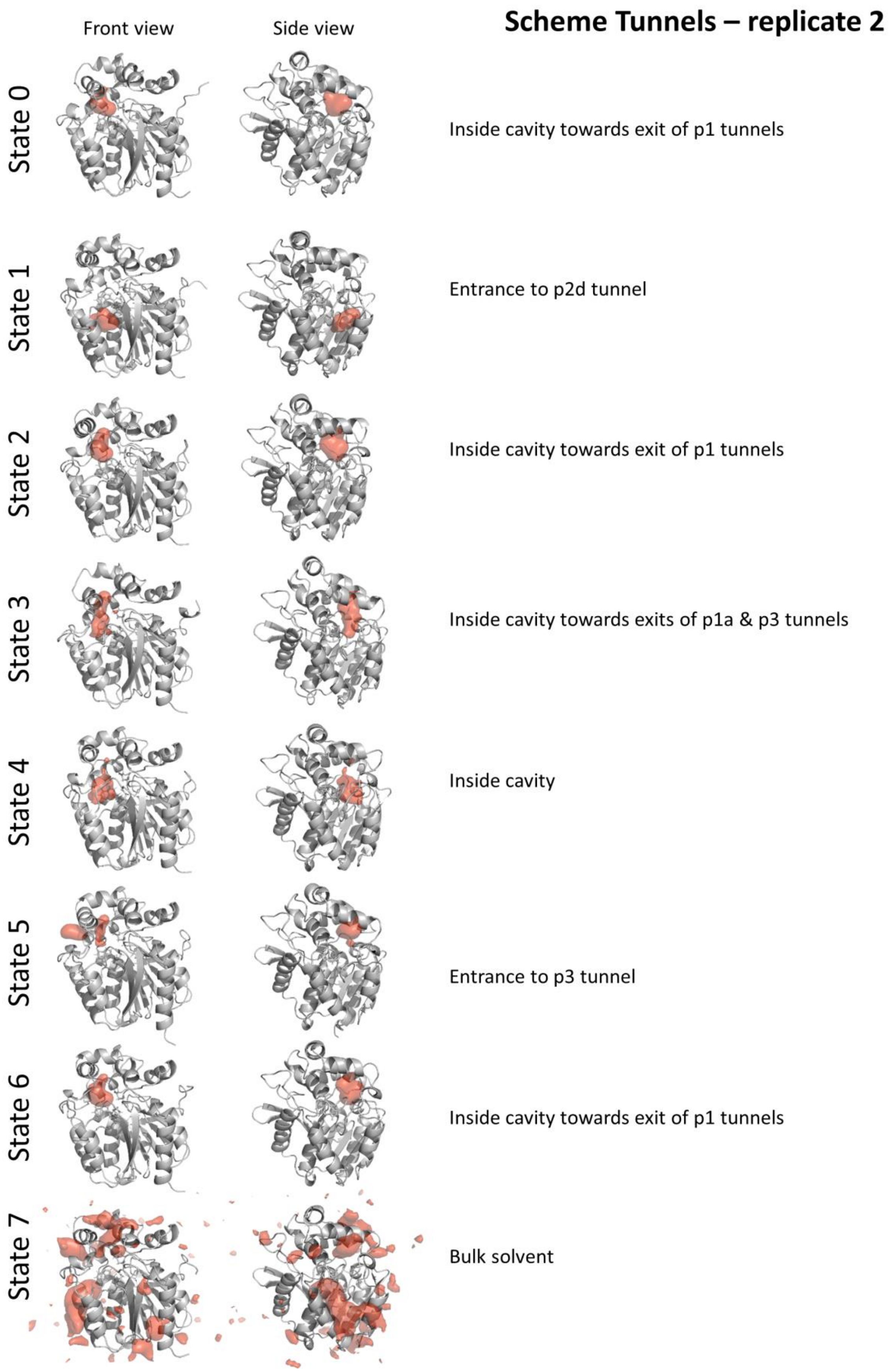


**Figure S24. Overview metastable states derived from MSM of *Tunnels* scheme replicate 2.** Protein structure is shown as a gray cartoon while the region occupied by DBE molecule in 20 % (1 % for bulk solvent state) of 1000 structures representing given metastable state is shown as red surface.

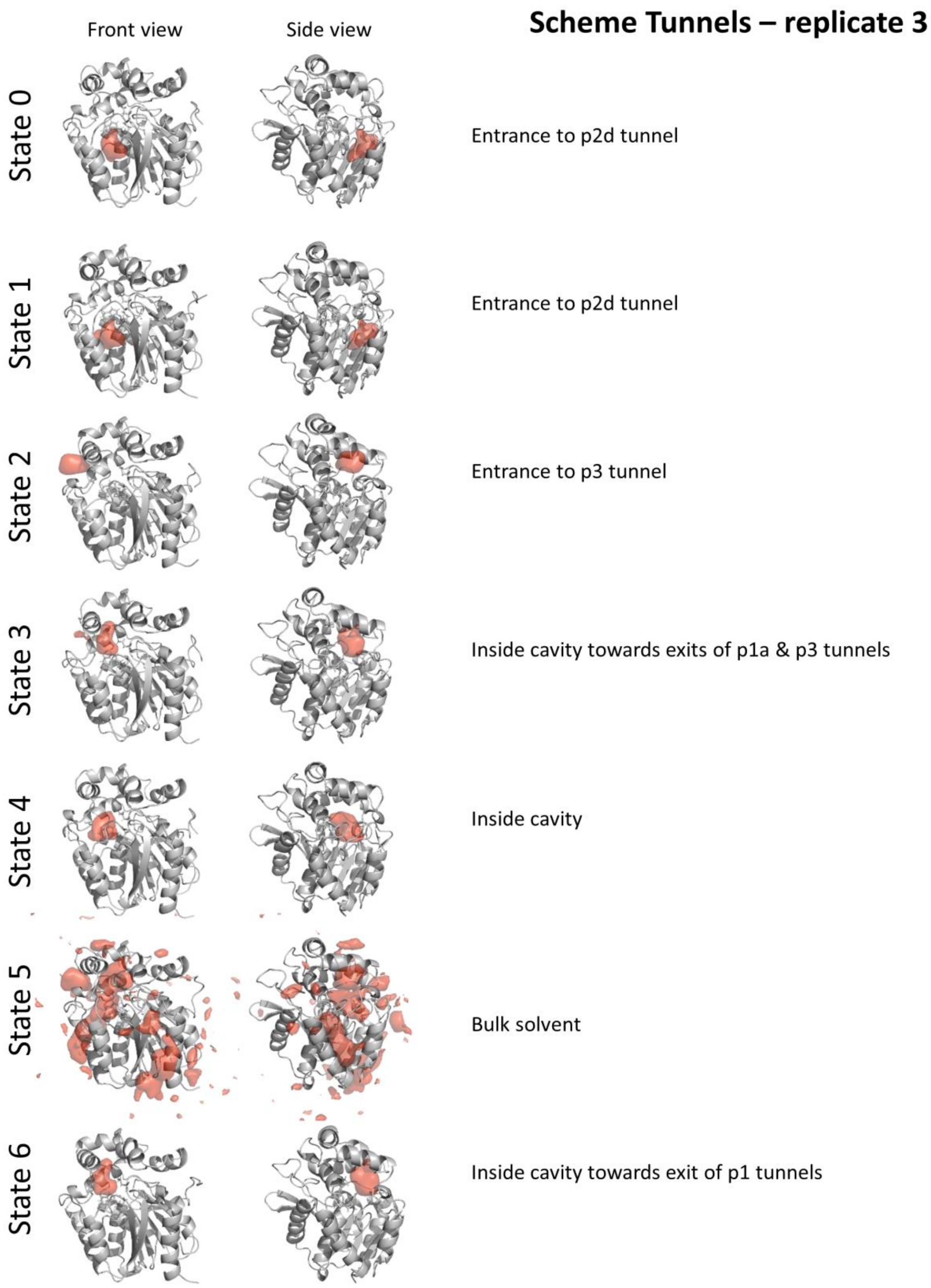


**Figure S25. Overview metastable states derived from MSM of *Tunnels* scheme replicate 3.** Protein structure is shown as a gray cartoon while the region occupied by DBE molecule in 20 % (1 % for bulk solvent state) of 1000 structures representing given metastable state is shown as red surface.

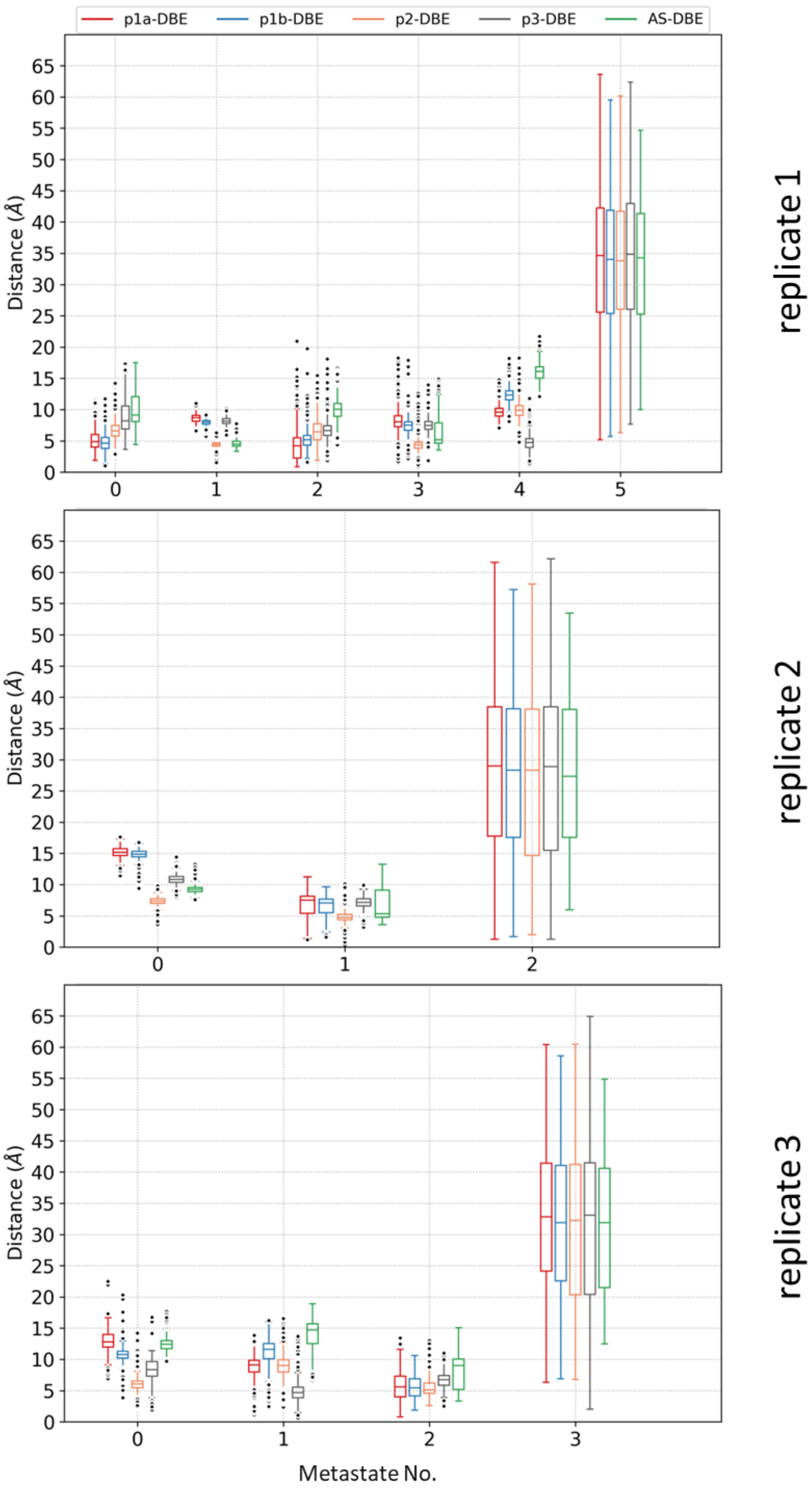


**Figure S26. Characteristic distances of representative structures of metastable states from the *Cavity* scheme.**

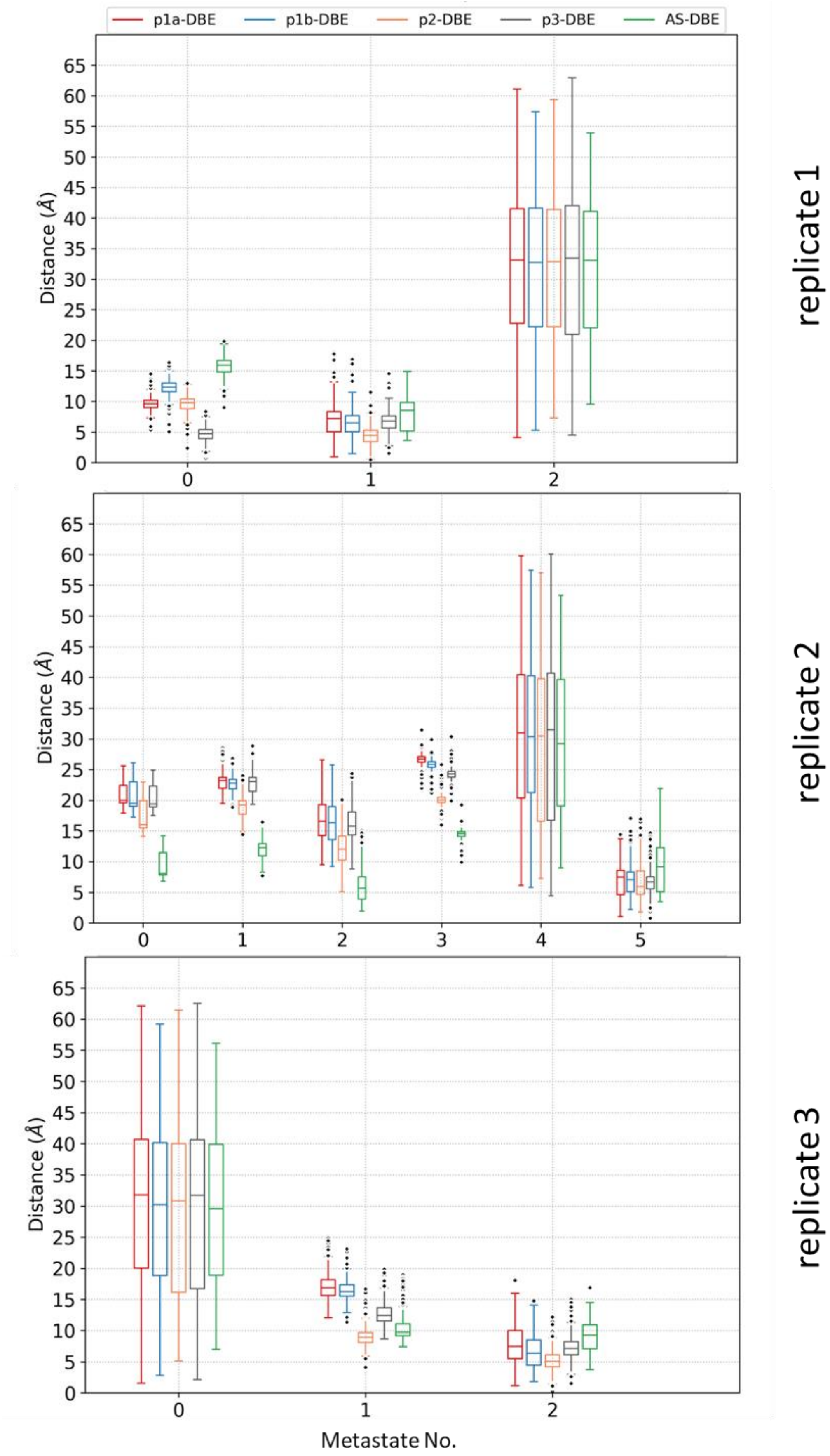


**Figure S27. Characteristic distances of representative structures of metastable states from the *Cavity&Bulk* scheme.**

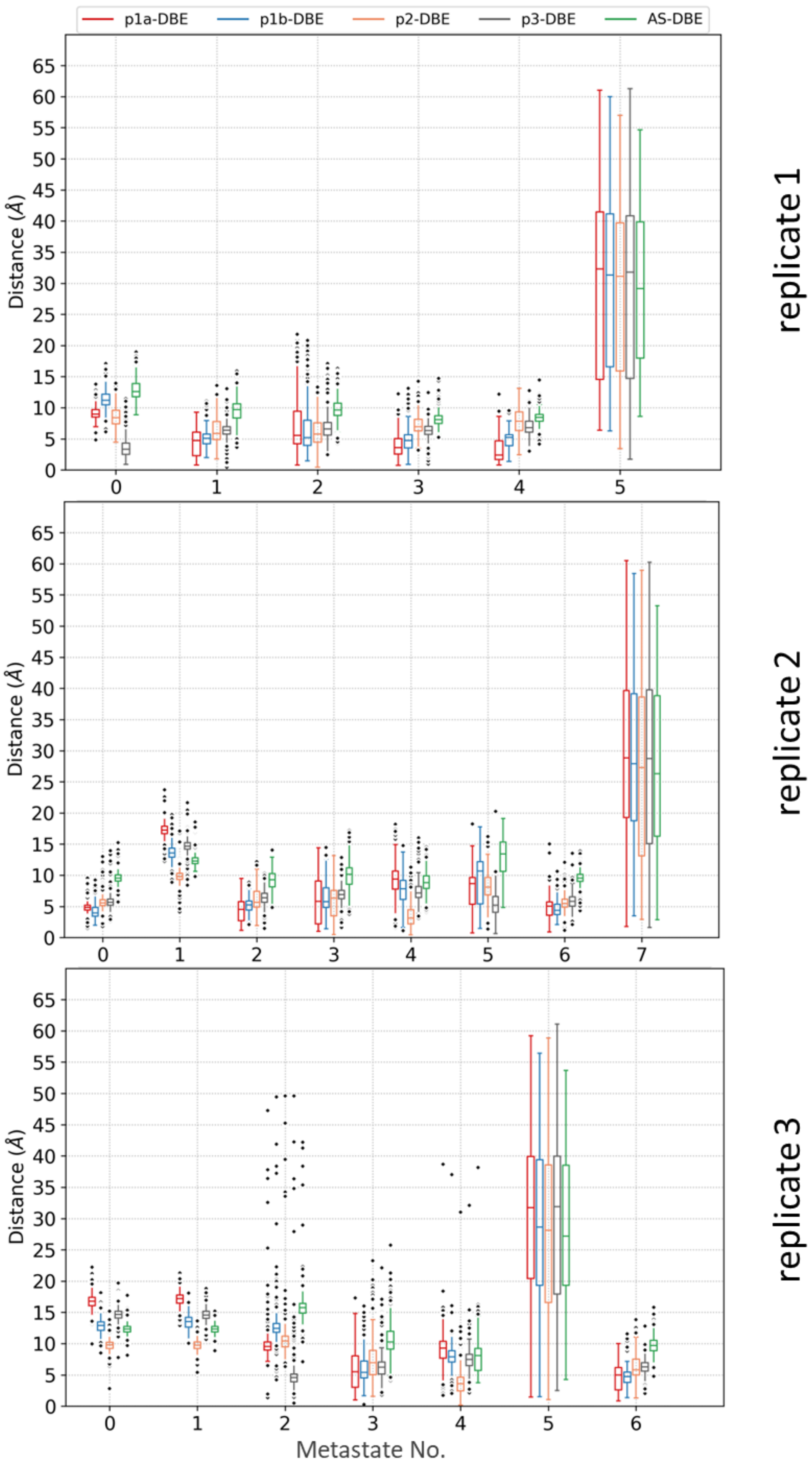


**Figure S28. Characteristic distances of representative structure of metastable states from the *Tunnels* scheme.**

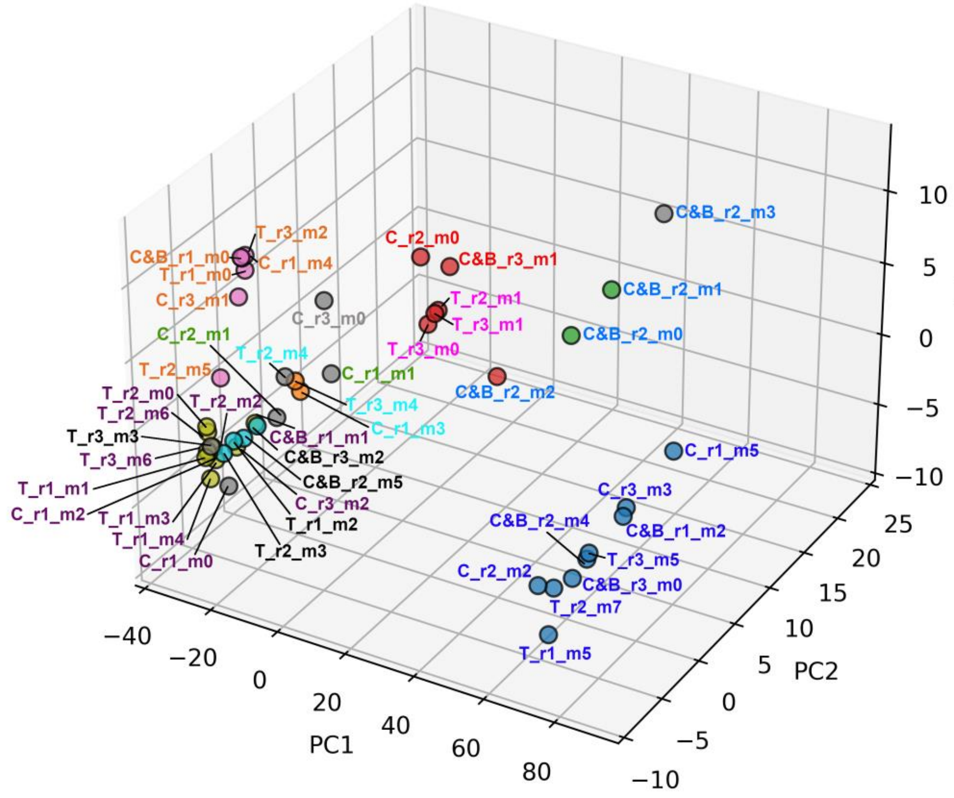


| ULS1 | ULS2 | ULS3 | ULS4 | ULS5 | ULS6 | ULS7 | ULS8 | ULS9 | ULS10 |
|---|---|---|---|---|---|---|---|---|---|
| C_r1_m5 | C_r1_m1 | T_r2_m4 | C_r1_m0 | C&B_r3_m2 | T_r3_m2 | C_r3_m0 | T_r2_m1 | C&B_r3_m1 | C&B_r2_m3 |
| C_r3_m3 | C_r2_m1 | T_r3_m4 | T_r1_m4 | C&B_r2_m5 | C_r1_m4 | | T_r3_m1 | C_r2_m0 | C&B_r2_m1 |
| C&B_r1_m2 | | C_r1_m3 | T_r1_m3 | T_r1_m2 | C&B_r1_m0 | | T_r3_m0 | | C&B_r2_m0 |
| C&B_r2_m4 | | | T_r2_m0 | T_r2_m3 | T_r1_m0 | | | | C&B_r2_m2 |
| T_r3_m5 | | | T_r2_m6 | T_r3_m3 | C_r3_m1 | | | | |
| C&B_r3_m0 | | | T_r2_m2 | | T_r2_m5 | | | | |
| T_r2_m7 | | | C&B_r1_m1 | | | | | | |
| C_r2_m2 | | | C_r3_m2 | | | | | | |
| T_r1_m5 | | | T_r3_m6 | | | | | | |
| | | | T_r1_m1 | | | | | | |
| | | | C_r1_m2 | | | | | | |

**Figure S29. Clustering of metastable states from all MSM analyses into unified ligand states (ULS).** The location of individual metastable states (Figures S17-S25) according to their fingerprints of characteristic distances (Figures S26-S28) in space formed by three principal components (PC) from the PC analysis. The disks are colored based on the cluster membership from the HDBSCAN method, with the gray disk corresponding to cluster composed of outliers. The metastable states are labeled according to the seeding scheme (C-*Cavity*, C&B-*Cavity&Bulk*, and T-*Tunnels*) followed by replicate (r1-3) and the metastable state number (n0-7) and colored by their assignment to the respective ULS.

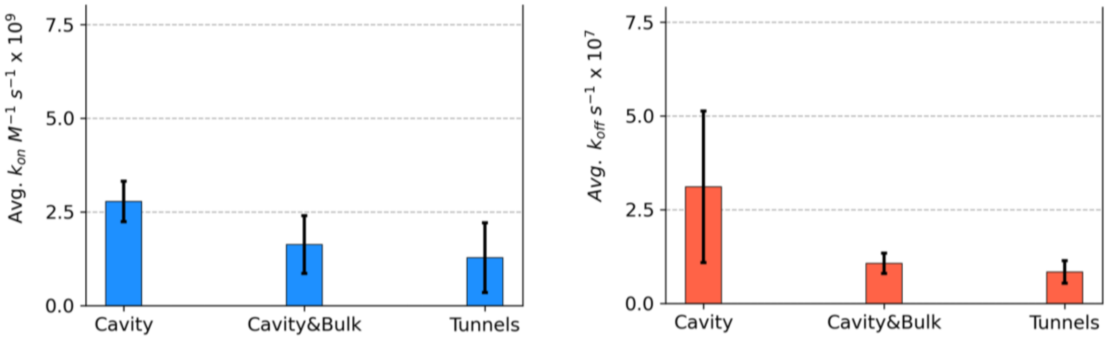


**Figure S30. Association and dissociation rates derived from MSMs.** The data represents mean±stdev from the three replicates.

**Table S3. Number of substrate migration events obtained from TransportTools analyses per schemes and replicates.**

| Schemes | Replicates | Total events | p1a | p1b | p2 | p3 | Unassigned events |
|---|---|---|---|---|---|---|---|
| Cavity | 1 | 11 | 0 | 0 | 0 | 0 | 11 |
| | 2 | 40 | 11 | 14 | 1 | 7 | 7 |
| | 3 | 7 | 0 | 0 | 0 | 0 | 7 |
| Cavity&Bulk | 1 | 5 | 0 | 0 | 0 | 0 | 5 |
| | 2 | 1 | 0 | 0 | 0 | 0 | 1 |
| | 3 | 3 | 0 | 0 | 0 | 1 | 2 |
| Tunnels | 1 | 6 | 0 | 0 | 0 | 1 | 5 |
| | 2 | 6 | 0 | 1 | 0 | 0 | 5 |
| | 3 | 10 | 0 | 0 | 0 | 0 | 10 |

**Table S4. Per scheme and tunnel counts and detailed statistics of tunnels' utilization by DBE ligand molecule.**

| Tunnel | rep1 | rep2 | rep3 | sum/tunnel | avg/tunnel | std/tunnel |
|---|---|---|---|---|---|---|
| **Bulk** | | | | | | |
| p1a | 358 | 17 | 156 | 531 | 177.0 | 171.5 |
| p1b | 527 | 4 | 293 | 824 | 274.7 | 262.0 |
| p2 | 614 | 15 | 404 | 1033 | 344.3 | 303.9 |
| p3 | 216 | 8 | 81 | 305 | 101.7 | 105.5 |
| unknown | 13 | 4 | 11 | 28 | 9.3 | 4.7 |
| mixed | 92 | 1 | 61 | 154 | 51.3 | 46.3 |
| sum/rep | 1820 | 49 | 1006 | | | |
| sum/scheme | 2875 | | | | | |
| avg/scheme | 958.3 | | | | | |
| std/scheme | 886.5 | | | | | |
| **Cavity** | | | | | | |
| p1a | 626 | 231 | 418 | 1275 | 425.0 | 197.6 |
| p1b | 525 | 160 | 883 | 1568 | 522.7 | 361.5 |
| p2 | 739 | 251 | 1084 | 2074 | 691.3 | 418.5 |
| p3 | 261 | 62 | 320 | 643 | 214.3 | 135.2 |
| unknown | 40 | 10 | 39 | 89 | 29.7 | 17.0 |
| mixed | 95 | 20 | 165 | 280 | 93.3 | 72.5 |
| sum/rep | 2286 | 734 | 2909 | | | |
| sum/scheme | 5929 | | | | | |
| avg/scheme | 1976.3 | | | | | |
| std/scheme | 1120.1 | | | | | |
| **Cavity&Bulk** | | | | | | |
| p1a | 241 | 113 | 131 | 485 | 161.7 | 69.3 |
| p1b | 239 | 80 | 689 | 1008 | 336.0 | 315.9 |
| p2 | 700 | 115 | 970 | 1785 | 595.0 | 437.1 |
| p3 | 143 | 71 | 155 | 369 | 123.0 | 45.4 |
| unknown | 6 | 39 | 25 | 70 | 23.3 | 16.6 |
| mixed | 47 | 16 | 84 | 147 | 49.0 | 34.0 |
| sum/rep | 1376 | 434 | 2054 | | | |
| sum/scheme | 3864 | | | | | |
| avg/scheme | 1288.0 | | | | | |
| std/scheme | 813.6 | | | | | |
| **Tunnels** | | | | | | |
| p1a | 363 | 306 | 261 | 930 | 310.0 | 51.1 |
| p1b | 931 | 425 | 854 | 2210 | 736.7 | 272.6 |
| p2 | 1071 | 726 | 1391 | 3188 | 1062.7 | 332.6 |
| p3 | 280 | 134 | 199 | 613 | 204.3 | 73.1 |
| unknown | 35 | 12 | 52 | 99 | 33.0 | 20.1 |
| mixed | 129 | 77 | 211 | 417 | 139.0 | 67.6 |
| sum/rep | 2809 | 1680 | 2968 | | | |
| sum/scheme | 7457 | | | | | |
| avg/scheme | 2485.7 | | | | | |
| std/scheme | 702.2 | | | | | |
